# *pIGLET*: Safe harbor landing sites for reproducible and efficient transgenesis in zebrafish

**DOI:** 10.1101/2023.12.08.570868

**Authors:** Robert L. Lalonde, Harrison H. Wells, Cassie L. Kemmler, Susan Nieuwenhuize, Raymundo Lerma, Alexa Burger, Christian Mosimann

## Abstract

Standard methods for transgenesis in zebrafish depend on random transgene integration into the genome followed by resource-intensive screening and validation. Targeted vector integration into validated genomic loci using phiC31 integrase-based *attP*/*attB* recombination has transformed mouse and *Drosophila* transgenesis. However, while the phiC31 system functions in zebrafish, validated loci carrying *attP*-based landing or safe harbor sites suitable for universal transgenesis applications in zebrafish have not been established. Here, using CRISPR-Cas9, we converted two well-validated single insertion *Tol2*-based zebrafish transgenes with long-standing genetic stability into two *attP* landing sites, called *phiC31 Integrase Genomic Loci Engineered for Transgenesis* (*pIGLET*). Generating fluorescent reporters, *loxP*-based Switch lines, CreERT2 drivers, and gene-regulatory variant reporters in the *pIGLET14a* and *pIGLET24b* landing site alleles, we document their suitability for transgenesis applications across cell types and developmental stages. For both landing sites, we routinely achieve 25-50% germline transmission of targeted transgene integrations, drastically reducing the number of required animals and necessary resources to generate individual transgenic lines. We document that phiC31 integrase-based transgenesis into *pIGLET14a* and *pIGLET24b* reproducibly results in representative reporter expression patterns in injected F0 zebrafish embryos suitable for enhancer discovery and qualitative and quantitative comparison of gene-regulatory element variants. Taken together, our new phiC31 integrase-based transgene landing sites establish reproducible, targeted zebrafish transgenesis for numerous applications while greatly reducing the workload of generating new transgenic zebrafish lines.

**SHORT ABSTRACT:** Targeted transgenesis into pre-established, “safe harbor,” landing sites remains a missing technique in zebrafish. Here, we established *phiC31 Integrase Genomic Loci Engineered for Transgenesis (pIGLET*) by CRISPR-Cas9-based conversion of two previously validated *Tol2* transgenes into *attP* sites. phiC31-mediated transgenesis into our *pIGLET14a* and *pIGLET24b attP* landing sites results in 25-50% germline transmission efficiency and quantifiable transgene activity. Our landing sites are suitable for reproducible, routine zebrafish transgenesis for diverse applications.

## INTRODUCTION

Transgenesis, the introduction of engineered DNA elements into the genome, is a key technique for any genetic model system. In zebrafish, transgenesis has resulted in invaluable fluorescent reporters and genetic modifier strains to study mechanisms of development, human disease, and physiology. However, routine transgenesis in zebrafish with random DNA integration via *Tol2* transposons or I-SceI Meganuclease presents considerable challenges: the creation of high-quality, single-copy transgene integrations with reproducible activity remains time-, labor-, and resource-intensive and faces unpredictable variability due to integration position effects that influence transgene activity^1–8^. While CRISPR-Cas9-mediated knock-in has greatly advanced the possibilities to generate targeted transgene integrations, success rates remain locus-dependent and variable^9–11^. Transgenesis into well-validated “safe harbor” sites for reproducible, routine transgenesis into predictable loci remains a technical hurdle in zebrafish, yet has the potential to positively transform the throughput, accessibility, and reproducibility of a key method in the model.

Adapted from the phiC31 bacteriophage, phiC31 integrase catalyzes the directional recombination of a vector-borne *attB* sequence into a genomic *attP* landing site that has been previously integrated by other transgenesis means^12–15^. However, the identification of safe harbor sites in model organism genomes has remained conceptually and practically challenging due to the uncertain influences of genomic environment (position relative to genes, regulatory elements, repetitive sequences, open chromatin regions, etc.) on individual transgene integrations^16,17^. Targeted vector integration using phiC31 integrase-mediated *attP*/*attB* recombination has transformed mouse and *Drosophila* transgenesis after well-defined, validated *attP* landing sites generated by random integration became accessible to the community^18–21^. While phiC31 functions in zebrafish and has been applied for regulatory element testing with fluorescent reporters following *Tol2*-based random transgenesis of *attP* sites^22–27^, universal *attP*-based landing sites that have been validated as potential safe harbors for a wide variety of transgene applications remain missing.

Here, converting two well-established *Tol2*-based transgenic integrations, we generated, validated, and applied two *attP* landing sites for key transgenesis applications using phiC31 integrase in zebrafish. We demonstrate their utility in reproducibly generating high-quality zebrafish transgenics, including *loxP*-based reporters that are overly sensitive to position effects. Accessible with standard methods applied in the field, our new landing sites enable qualitative and quantitative transgene applications, facilitate reproducible reagent generation and sharing, as well as streamline the transgenesis workflow while reducing required effort and animal numbers. Beyond two widely applicable landing sites for community use, our work proposes that conversion of high-quality *Tol2* transgenes into *attP* landing sites has the potential to drastically increase the number of transgenesis landing sites available in the field.

## RESULTS

### Converting two functional *loxP* transgenes into *attP* landing sites

Due to random integration, transgenes generated with the *Tol2* transposon system or I-SceI (fluorescent reporters, CreERT2-drivers, *UAS*-effectors, etc.) are susceptible to position effects, requiring labor- and resource-intensive screening to identify transgenic founders with accurate, reproducible expression^1,4^. A key tool for genetic lineage labeling, *loxP*-based reporter transgenes that recombine upon Cre recombinase activity, are also notably susceptible to position effects in zebrafish and challenging to generate^4^. As they retain access for Cre recombinase action, we reasoned that well-recombining and -expressing *loxP* reporter transgenes present suitable integration sites for transgenesis. We previously generated and characterized the *loxP*-based Cre reporter strains *ubi:loxP-EGFP-loxP_mCherry* (*ubi:Switch*) and *hsp70l:loxP-STOP-loxP_EGFP* (*hsp70l:Switch*) as *Tol2* transgene integrations with efficient recombination and expression^17,28,29^. Both lines i) have been in use for generations in numerous labs, ii) feature dependable, reproducible *loxP* recombination upon Cre activity and iii) are homozygous viable without appreciable phenotypic consequences. We therefore performed CRISPR-Cas9-based knock-in of phiC31-recognized *attP* landing sites to replace the *Tol2* transgenes in these strains to generate alleles entitled *<u>p</u>hiC31Integrase <u>G</u>enomic <u>L</u>oci <u>E</u>ngineered for <u>T</u>ransgenesis* (*pIGLET*) (**Fig. S1**).

Our previous mapping of the genomic locations of both *ubi:Switch* and *hsp70l:Switch* had revealed that both had integrated into repetitive sequences^17^, barring us from designing locus-specific CRISPR-Cas9 targeting strategies. We therefore devised a two-step process to generate *attP* landing sites in each locus that is applicable to any previously generated *Tol2* transgene (**Fig. S1**): *Tol2*-based transgenes are flanked by unique 5’ and 3’ *Tol2* repeats, which we targeted using individual sgRNAs for Cas9 RNP-mediated transgene excision in homozygous transgenic embryos (**Fig. S1A**). Screening for transgene-negative F1 embryos for both lines (see **Methods**), we identified clean deletions of the integrated transgenes (1/20 F0-injected embryos) that left a unique *5’-3’ Tol2* repeat remnant sequence in the genome (Step 1) (**Fig. S1A-C,F,G**). We then targeted this sequence in embryos from heterozygous in-crosses using a single sgRNA and coinjected a single-stranded oligonucleotide (ssODN) containing an 84 bp-spanning *attP* sequence flanked by asymmetric homology arms of the remnant *Tol2* repeats (**Fig. S1D**, see Methods for details). We identified *attP* integrations in both the *ubi:Switch* (1/18 F0-injected embryos) and *hsp70l:Switch* (1/35 F0-injected embryos) and designated these new alleles *pIGLET14a* and *pIGLET24b*, respectively (Step 2) (**Fig. S1E,H,I)**. Both *attP* integrations are homozygous viable and detectable with simple PCR genotyping for line maintenance (**see Methods**).

To standardize nomenclature, we propose to name *pIGLET* landing sites with chromosome number and a consecutive letter index: *pIGLET14a^co2001^* and *pIGLET24b^co2002^*. We further propose to refer to transgene insertions into individual landing sites as *Tg(pIGLET14a.transgene)* or *Tg(pIGLET24b.transgene)*, shortened as *p14a.transgene* and *p24b.transgene* to distinguish loci in the text (e.g., *p14a.drl:EGFP* and *p24b.drl:EGFP*).

### *pIGLET14a* and *pIGLET24b* are highly efficient landing sites

We next sought to test the *pIGLET14a* and *pIGLET24b* landing sites for phiC31 integrase-mediated transgenesis. A typical workflow to implement the *pIGLET* system for zebrafish transgenesis is described in **Fig. 1A,B**, including a schematic depicting targeted transgene integration into the landing sites and validation by PCR and sequencing. Importantly, PCR-based genotyping of *attP*/*attB* recombination in *pIGLET14a* and *pIGLET24b* landing sites for quality control and validation of phiC31 *integrase* mRNA can be performed in stable transgenic lines and F0-injected embryos, with documented recombination within the first 24 hpf in 100% of injected embryos with observable transgene expression (see **Supplementary Methods**). This simple quality control step provides confidence in the methods and reagents used, in a manner comparable to the now universal F0-testing of sgRNA functionality for CRISPR-Cas9 mutagenesis^30^. In addition to our previously established *attB-*containing *pDEST* backbone vectors (*pCM268*, no transgenesis marker; *pCM327*, *cryaa:Venus*)^22^, we generated six additional backbone vectors for different cloning approaches: *pRL055 pDESTattB_cryaa:Venus* (inverted), *pRL56 pDESTattB_exorh:EGFP*, *pCK122 pDESTattB_exorh:mCherry*, *pCK123 pDESTattB_exorh:mCerulean, pRL092 pattB_cryaa:Venus_MCS, and pRL093 pattB_exorh:EGFP_MCS* (for details, see **Table 1** and Methods). While *cryaa:Venus* provides an eye lens-specific transgenesis marker, the *exorh* element provides pineal gland-specific reporter expression for simple transgene screening (**Table 1**)^31^. The *attB* site-containing *pDEST* vectors *pCM268, pRL055, pRL056, pCK122,* and *pCK123* are designed as backbones to generate expression plasmids using the Multisite Gateway system in combination with *p5E, pME,* and *p3E* plasmids^2,31^. Alternatively, the *attB* site vectors *pRL092* and *pRL093* enable restriction enzyme- or PCR-based cloning and contain a multiple cloning site (MCS) with eight available unique restriction sites (*NsiI, EcoRV, NheI, NdeI, SacI, BstEII, StuI, AgeI*) (**Table 1**). All vector maps and details of genomic locations of the landing sites are documented in the **Supplementary Data** file (.zip).

**Figure 1.**
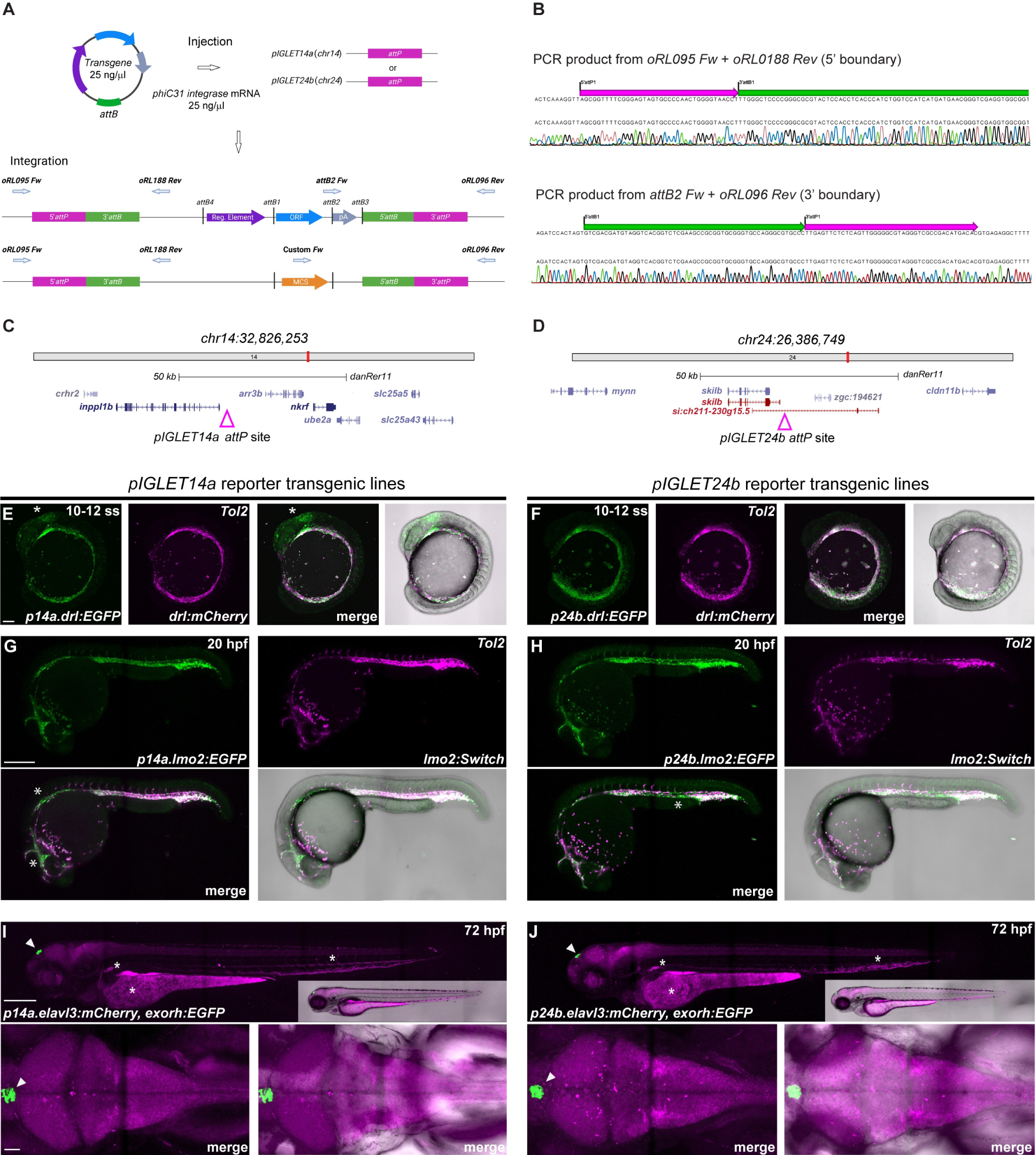
phiC31 integrase-mediated transgenesis into *pIGLET14a* and *pIGLET24b* landing sites in zebrafish reproducibly results in high-quality transgenic lines. (**A, B**) Schematic of *pIGLET* system workflow (injection, integration, sequencing). Expression vectors are assembled in *attB*-containing *pDEST* backbones and then co-injected with phiC31 *integrase* mRNA into *pIGLET14a* or *pIGLET24b* zebrafish (**A**). Following transgene integration with either, 5’ and 3’ transgene boundaries can be confirmed via PCR and sequencing (**B**). Integration with *pDESTattB* vectors (*pCM268, pRL055, pRL056, pCK122, pCK123*) and *pattB_MCS* vectors (*pRL092, pRL093*) are depicted (**B**). Genomic locations of *pIGLET14a and pIGLET24b* landing sites are depicted in **C** and **D** and annotated .gbk sequence files of both landing site integrations are included as Supplementary Data. (**E-J**) Validation of *pIGLET14a* and *pIGLET24b* loci for three reporter transgenes*: drl:EGFP, lmo2:EGFP,cryaa:Venus* and *elavl3:mCherry,exorh:EGFP*. (**E, F**) Crossed to *Tol2*-based *drl:mCherry* (magenta fluorescence), the *pIGLET* transgenes *p14a.drl:EGFP* and *p24b.drl:EGFP* show analogous reporter activity in the lateral plate mesoderm at 10-12 ss (green fluorescence), consistent with well-established *Tol2*-based *drl* reporter transgenics. Note faint ectopic brain expression in *p14a.drl:EGFP* at 10-12 ss (**E**, white asterisk) that is common with *Tol2*-based *drl* reporter transgenics with consistent and strong expression levels. (**G, H**) Crossed to *Tol2*-based *lmo2:Switch (*dsRed2, magenta), *p14a.lmo2:EGFP* and *p24b.lmo2:EGFP* show analogous, overlapping reporter activity in the endothelial lineages at 20 hpf. *EGFP* expression is more consistent/complete compared to *dsRED2* expression observed in *lmo2:Switch* (**G, H**, white asterisk). (**I, J**) *p14a.elavl3:mCherry* and *p24b.elavl3:mCherry* both show complete, consistent reporter expression in the central nervous system (brain, spinal cord) at 72 hpf, in line with previously described *elavl3* transgenics. Lateral views and dorsal views showing the *in cis exorh:EGFP* transgenesis marker (white arrowhead). Note common autofluorescence in blood, kidney, and yolk (**I**, **J**, white asterisk). Scale bars: C, D = 100 μm; E, F = 250 μm; G, H = 250 μm, 50 μm.

**Table 1.**
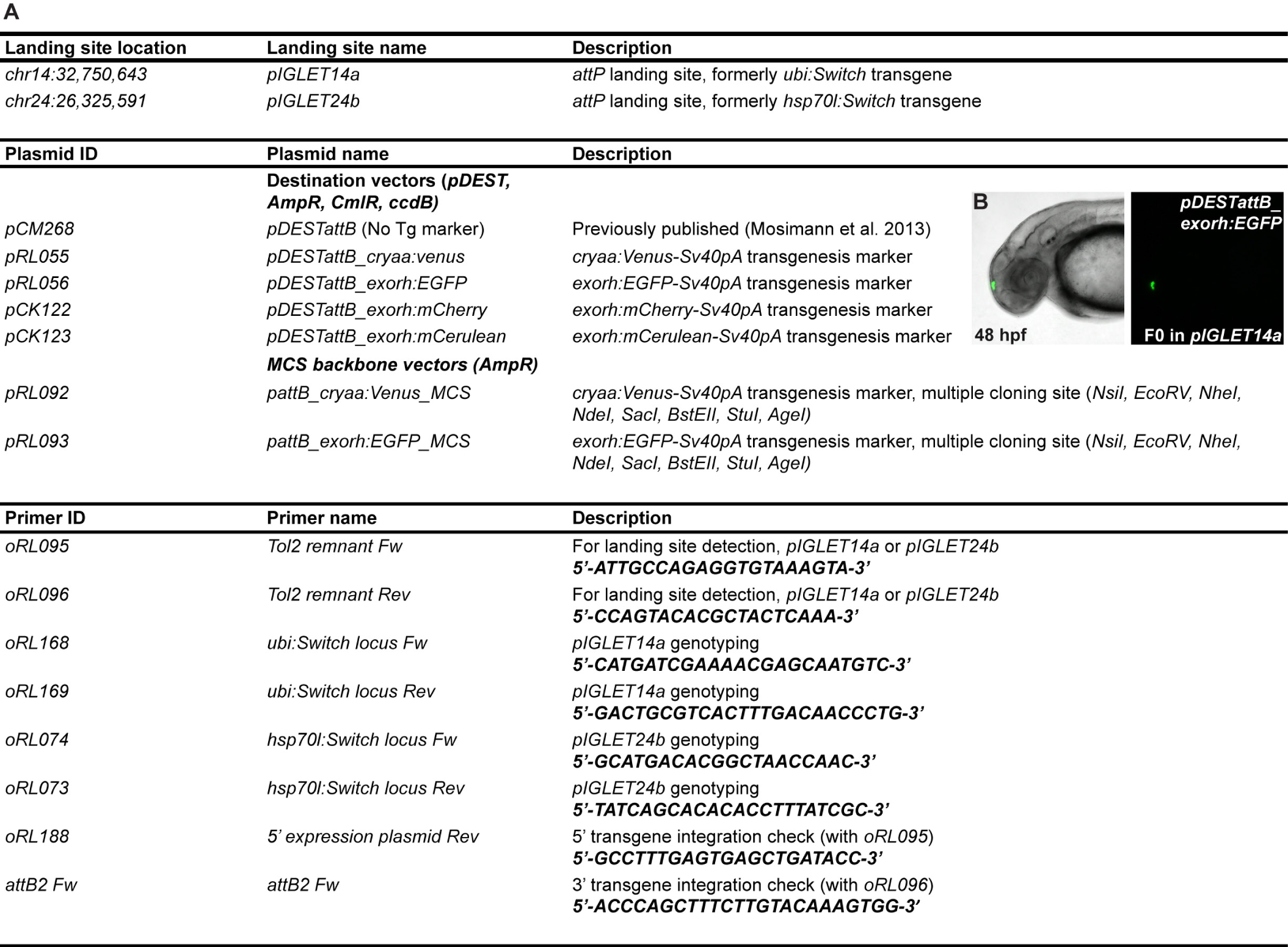
*pIGLET* reagents. Table of landing site information, *pDEST* plasmids, and primers for *pIGLET* system implementation (**A**). F0-injection of newly created *attB*-containing *pDEST* vectors with *exorh:EGFP* transgenesis marker (**B**). See also Supplementary Data file (.zip) for vector maps and genomic loci (GenBank format).

Using phiC31 integrase-mediated transgenesis, we next sought to integrate different *attB*-based transgenic reporter and effector vectors into *pIGLET14a* and *pIGLET24b* (**Table. S1, Table. S2**). Accessible with standard methods applied in the field, phiC31 integrase-mediated transgenesis is performed by co-injection of an *attB* sequence-containing transgenesis vector together with mRNA encoding for the phiC31 integrase into one-cell stage zebrafish embryos, at comparable concentrations to those used in Tol2-mediated transgenesis (e.g., 25 ng/µl vector, 25 ng/µl *integrase* mRNA). A key difference to other transgenesis approaches is the need to inject into zebrafish that carry an *attP* landing site, such as *pIGLET14a* and *pIGLET24b.* Consequently, we recommend injection into homozygous landing site embryos to provide optimal transgenesis efficiency. In the subsequent paragraphs, newly created transgenic lines will have *p14a* and *p24b* designations to reflect their *pIGLET* integration locus (e.g., *p14a.drl:EGFP*, *p24b.drl:EGFP*). Transgene integration in the *pIGLET14a* and *pIGLET24b* loci is highly efficient, routinely achieving 25-50% germline transmission from F0-injected zebrafish. A complete record of germline screening statistics for all lines recovered in the *pIGLET14a* and *pIGLET24b* loci is compiled in **Table S2**.

*Tol2*-based fluorescent reporters harnessing the 6.35 kb *draculin* (*drl*) regulatory elements mark the emerging lateral plate mesoderm (LPM) and subsequently refine to cardiovascular and hematopoietic lineages^32–34^ (**Fig. 1E,F; Fig. S2A,B, Fig. S3A,B**). To re-derive *drl*-based reporters in each *pIGLET* landing site, we re-cloned the well-established reporter *drl:EGFP* into an *attB* site-carrying vector backbone and injected with phiC31 integrase-encoding mRNA into heterozygous in-crossed *pIGLET14a* and *pIGLET24b* offspring. Subsequent screening of potential founders resulted in over 50% of injected zebrafish with germline-transmitting *drl:EGFP* in the corresponding landing site (**Table. S2**). For both landing sites, the resulting expression pattern and dynamics were comparable to the *Tol2*-based, *drl:mCherry* reporter transgenic line that was previously selected for faithful expression identical to *Tol2*-based *drl:EGFP* integrations (**Fig. 1E,F; Fig. S2A,B, Fig. S3A,B**)^33^.

The regulatory elements of *lmo2* mark emerging endothelial and hematopoietic cell types^35^ (**Fig. 1G,H; Fig. S2C,D**); however, *lmo2*-based fluorescent reporter constructs are highly susceptible to position effects, requiring screening of several independent random insertions to isolate faithful reporter activity^35,36^. Germline-transmitted integrations of *p14a.lmo2:EGFP* and *p24b.lmo2:EGFP* showed labelling of vasculature and blood, and more uniform expression compared to the previously described, *Tol2*-based *lmo2:Switch* transgenic line (*lmo2:loxP-dsRED2-loxP_EGFP*) (**Fig. 1G,H; Fig. S2C,D**)^36^.

To investigate landing site functionality for non-mesodermal lineages, we created transgenic reporter lines based on the regulatory elements of *elavl3* (*HuC*), a pan-neuronal marker^37^. Using the previously described 8.7 kb regulatory region^38,39^, we created *elavl3:mCherry, exorh:EGFP* transgenic lines in both the *pIGLET14a* and *pIGLET24b* loci. *p14a.elavl3:mCherry* and *p24b.elavl3:mCherry* transgenes show expression in the brain and spinal cord comparable to previously described transgenic lines^38,39^ (**Fig. 1I,J**).

Examination of adult zebrafish confirmed that expression of transgenesis markers *cryaa:Venus,* and *exorh:EGFP* are maintained throughout development in both *pIGLET14a* and *pIGLET24b* landing sites (**Fig. S4A-D**). PCR-based genotyping of the recombined *attP*/*attB* sites confirmed precise transgene integrations of all characterized strains (see **Supplementary Methods, Fig. 1B, Table S2**). Together, these results underline the safe harbor potential of *pIGLET14a* and *pIGLET24b* for reproducible, high-quality transgenesis of fluorescent reporters with complex developmental expression patterns.

### *pIGLET14a* and *pIGLET24b* allow predictable generation of Cre/*lox* transgenes

To test whether our landing sites are suitable to generate functional CreERT2 driver transgenes, we created *drl:creERT2* transgenic lines in both the *pIGLET14a* and *pIGLET24b* loci and compared their activity to the well-characterized *Tol2-*based *drl:creERT2* transgene^32,40–43^. CreERT2 driver transgenes are also highly sensitive to position effects, with the highly efficient, *Tol2-*based *drl:creERT2* transgenic line requiring screening of ten independent founders^32^. By performing Cre/*loxP* lineage trace experiments with *hsp70l:Switch*, we compared the dynamics of the *drl* regulatory element in resolving complex LPM-based lineages (**Fig. 2A**). As previously described, shield stage 4-OHT induction using *Tol2-*based *drl:creERT2* results in complete LPM lineage labeling (heart, pectoral fin, endothelial cells, blood, kidney, pharyngeal arch musculature) and a subset of paraxial lineage labeling (skeletal muscle, median fin fibroblasts) (**Fig. 2A**). A 4-OHT induction series demonstrates that by 12 ss (somite stage), only LPM lineages are labeled using *drl:creERT2*, albeit with notable mosaicism (**Fig. 2A**). This dynamic, induction time-point dependent lineage labeling is recapitulated in the newly generated *p14a.drl:creERT2* and *p24b.drl:creERT2* lines in the *pIGLET14a* and *pIGLET24b* loci (**Fig. 2B, Fig. S5A**). Notably, *pIGLET* landing sites now enable integration of CreERT2, Gal4 drivers, and other effector transgenes at the same locus as their reporter transgene counterpart, providing confidence in the activity of the regulatory elements across transgene types.

**Figure 2.**
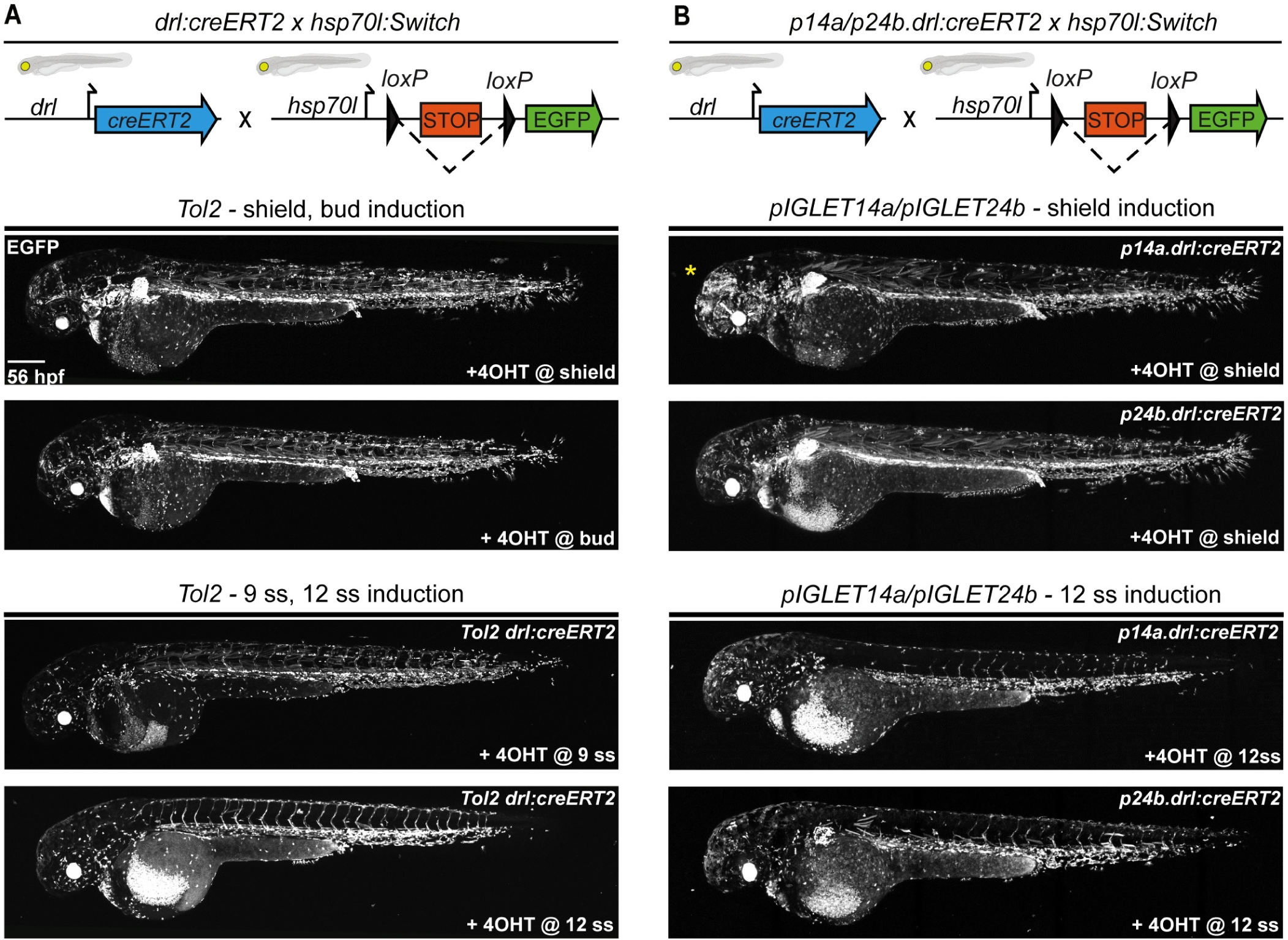
CreERT2 driver transgenes have predictable and consistent activity in *pIGLET14a and pIGLET24b*. Comparison of CreERT2-mediated recombination of *Tol2-*based *drl:creERT2* and *pIGLET*-based lines *p14a.drl:creERT2,* and *p24b.drl:creERT2* by crossing to the *loxP* reporter *hsp70l:Switch* and performing a 4-OHT induction series. Transgene and crossing schematics on top, lateral confocal views, anterior to the left. (**A, B**) Compared to the *Tol2*-based *drl:creERT2* transgenic line, *p14a.drl:creERT2* and *p24b.drl:creERT2* show accurate lineage contribution dynamics at 56 hpf when treated with 4-OHT at shield stage and 12 ss (**A, B**). Note with shield stage 4-OHT induction, *drl*-expressing cells contributed both to the lateral plate mesoderm lineages (heart, pectoral fin, endothelial cells, kidney, mesothelia, blood, pharyngeal arch musculature, macrophages) and a subset of paraxial mesoderm lineages (median fin fibroblasts, skeletal muscle). With 12 ss 4-OHT induction, lateral plate mesoderm lineages are specifically traced with more mosaic labeling of heart, pectoral fin, and kidneys (**A, B**). Consistent with our *p14a.drl:EGFP* reporter lines, some neuronal lineage labeling is observed with *p14a.drl:creERT2* when induced with 4-OHT at shield stage (Yellow asterisk) (**B**). Scale bars: A, B = 200 μm.

We next sought to test if *loxP*-based reporters for Cre activity (so-called *Switch* lines) are functional in our new landing sites. *loxP*-based *Switch* reporters have a high sensitivity to position effects and are therefore challenging to generate using random integrations, requiring resource-intensive screening efforts to isolate well-recombining *Switch* transgene integrations^1,4,17,28,29,44–47^. As the original *ubi:Switch* and *hsp70l:Switch* have high *loxP* recombination efficiency, we introduced a new *Switch* transgene termed *hsp70l:Switch2* (*hsp70l:loxP-STOP-loxP_mApple, cryaa:Venus*) that recombines from no fluorophore to mApple (red fluorescent), into each the two new *pIGLET* loci.

We tested the recombination efficiency of *hsp70l:Switch2* in both *pIGLET14a* and *pIGLET24b* with several previously published CreERT2-driver lines: *ubi:creERT2*^29^*, drl:creERT2*^32^, and *crestin:creERT2*^48^, each induced with 4-OHT and imaged at 56 hpf (**Fig. 3A,B**). When crossed to *ubi:creERT2* and *drl:creERT2*, *hsp70l:Switch2* displayed high levels of recombination and reporter expression upon heat-shock akin to the previously described *hsp70l:Switch*^17,28,49,50^ (**Fig. 2A**, **Fig. 3A,B**). *crestin* is expressed in early neural crest cells during early somitogenesis and becomes undetectable by 72 hpf, with *crestin:creERT2* enabling mosaic lineage-labeling of neural crest cells throughout development, instrumental for lineage studies and tracking of melanoma tumors^48,51–53^. Consistent with *hsp70l:Switch*-based lineage labeling (**Fig. S5D**) and previous observations using *ubi:Switch*^48^, we crossed *hsp70l:Switch2* to *crestin:creERT2* and observed mosaic recombination in neural crest cells (**Fig. 3A, B, Fig. S5B,C**). Taken together, our benchmarking establishes that *loxP*-based Switch reporters show faithful expression and sensitive, reproducible Cre-mediated recombination when integrated into the *pIGLET14a* and *pIGLET24b* landing sites. This property facilitates the predictable, efficient generation of new Cre-responsive *Switch* reporters and effector lines in zebrafish, overcoming a critical technical bottleneck in the field.

**Figure 3.**
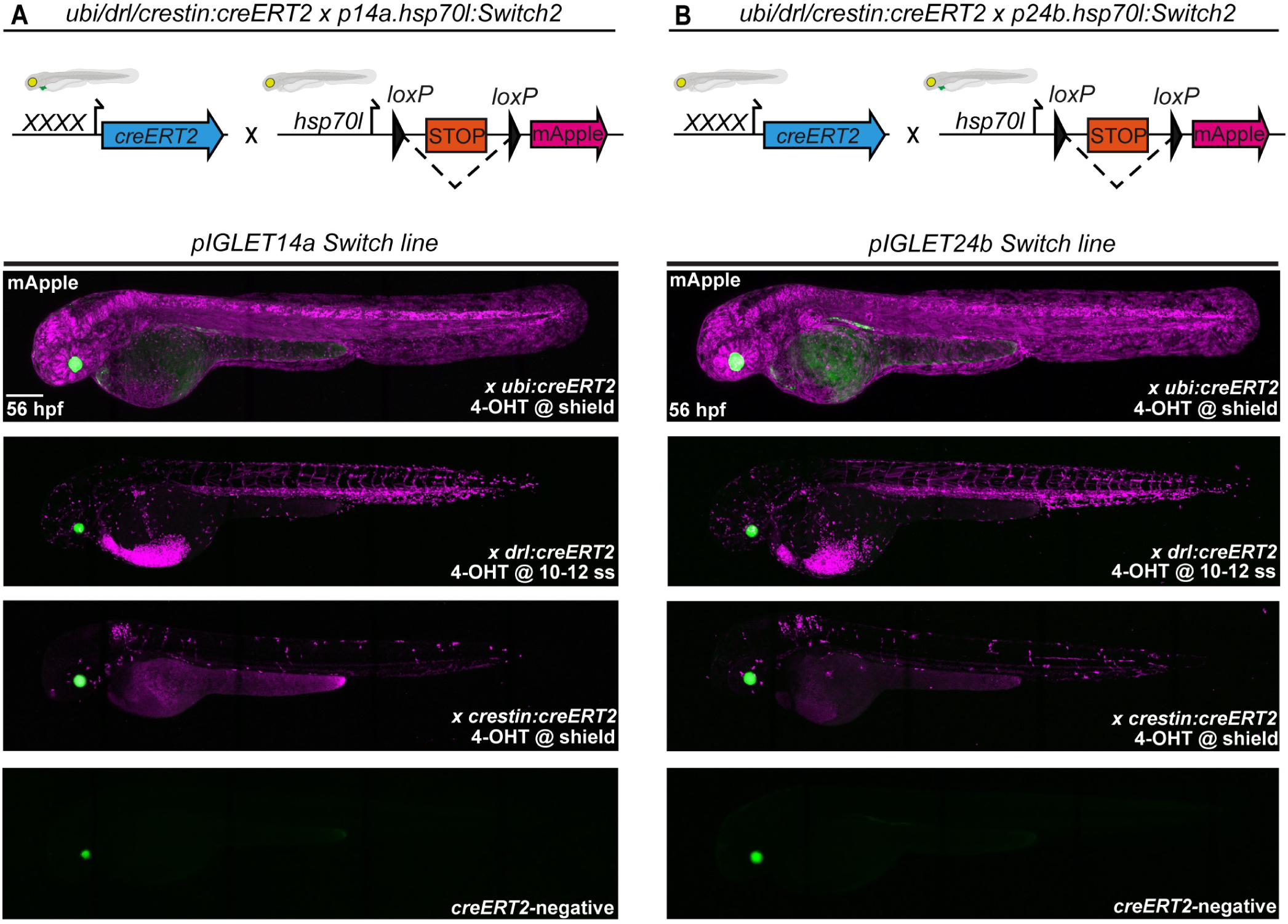
*loxP Switch* transgenes have predictable and consistent activity in *pIGLET14a* and *pIGLET24b*. Comparison of CreERT2-mediated recombination of *pIGLET*-based *loxP* reporter *hsp70l:Switch2*. Transgene and crossing schematics on top, lateral confocal views, anterior to the left. (**A, B**) Testing of *p14a.hsp70l:Switch2* and *p24b.hsp70l:Switch2* with three independent CreERT2-driver lines: *ubi:creERT2, drl:creERT2,* and *crestin:creERT2*. *p14a.hsp70l:Switch2* and *p24b.hsp70l:Switch2* drive strong *mApple* reporter expression following CreERT2-mediated recombination when crossed to ubiquitous and tissue-specific CreERT2-driver lines, treated with 4-OHT, and imaged at 56 hpf (**A, B**). Note the near complete *mApple* labeling of the embryo at 56 hpf when *p14a.hsp70l:Switch2* and *p24b.hsp70l:Switch2* are combined with *ubi:creERT2*, and accurate labeling of the lateral plate mesoderm or neural crest cells when used in combination with *drl:creERT2* or *crestin:creERT2,* respectively, akin to previous results using *Tol2*-based versions of either CreERT2 driver (**A, B**). Scale bars: A, B = 200 μm.

As previously documented for vector injection-based transgenesis in zebrafish by any method^1,22,26^, approximately 8-10% of germline-transmitted events result from random background integration caused by transgene vector that linearized at different locations in its sequence. We also uncovered such events using *pIGLET*-targeted phiC31 integrase-mediated transgenesis based on lower or variable transgene expression among reporter-expressing embryos (6/76 clutches analyzed, 7.89%) (**Table S2**). These events predominantly occurred with plasmid sizes of ∼14kb or greater (5/6 non-targeted integration clutches) (**Table S2**). PCR-based genotyping for the genomic *attP* integration or *attP*/*attB* recombination as well as germline segregation patterns of these transgenic insertions confirmed their non-targeted integration (**see Supplementary Methods**). Similar integrations have been well-documented in mice when targeting the *attP* sites in the *H11* locus, documenting background integrations as a universal caveat of vector-based transgene injections^18,21^. Nonetheless, our genotyping PCR to verify proper transgene integration into *attP* landing sites together with careful screening enable selection of correctly targeted transgenes into the *pIGLET14a* and *pIGLET24b* landing sites.

### Functional testing of enhancers using *pIGLET* landing sites

Targeted transgenesis into a known and robust genomic location opens the possibility of qualitative and quantitative reporter comparisons of gene-regulatory elements between wild type and variants such as polymorphisms, transcription factor (TF) binding site deletions, etc. (applications which have remained challenging with random transgenesis). We first sought to validate this approach using *pIGLET* by testing TF binding site deletion in the autoregulatory *Hs_I* notochord enhancer of the human *Brachyury*/*TBXT* gene that drives reporter activity in the notochord of zebrafish, axolotl, and mice (**Fig. 4A,B**)^54^. Removal of a Tbox motif in *Hs_I* results in loss of reporter expression in F0-injected zebrafish using *Tol2-*transgenesis constructs^54^. To recapitulate this result using the *pIGLET* system, we cloned the wild type and the Tbox deletion variant of enhancer *Hs_I* into *attB* site-carrying reporter vectors (*Hs_I:min-mCerulean,exorh:EGFP and Hs_IΔTbox:min-mCerulean,exorh:EGFP*) (**Fig. 4B-D**). To compare F0 notochord reporter mosaicism between *pIGLET* and *Tol2*-based transgenesis, we injected *Hs_I:min-mCerulean,exorh:EGFP* into *pIGLET14a* homozygous embryos and *Tol2*-based *Hs_I:min-mApple,exorh:EGFP*^54^ into wild type embryos in parallel and scored F0 expression based on percentage of notochord fluorescence coverage (**Fig. 4B-E, Fig S6A, B**). Injection of the Tbox deletion variant *Hs_IΔTbox:min-mCerulean, exorh:EGFP* into *pIGLET14a* resulted in complete loss of notochord expression, consistent with previous *Tol2-*based F0 results^54^ (**Fig. 4D**). Although both transgenesis methods achieved F0 embryos with 80-100% fluorescent coverage across notochords for the wild type *Hs_I* reporters (**Fig. 4B,C,E**), *pIGLET*-based injections yielded a significantly higher percentage of nearly complete notochord reporter activity among injected embryos (*Tol2: 4.72% vs. pIGLET*: 27.42%) (**Fig. 4E**). This result documents lower F0 mosaicism using the *pIGLET* system compared to *Tol2* and is consistent with our observations of F0 expression following injection of *drl:EGFP*, and *lmo2:EGFP* constructs into homozygous landing site embryos (90-95% transgene-positive in injected clutches, approximately 25% displaying strong expression) (**Fig. S7A-H**). Indicative of early genomic integration after injection, this additional feature of our two *pIGLET* lines potentially facilitates selection and screening of regulatory elements for subsequent germline transmission.

**Figure 4.**
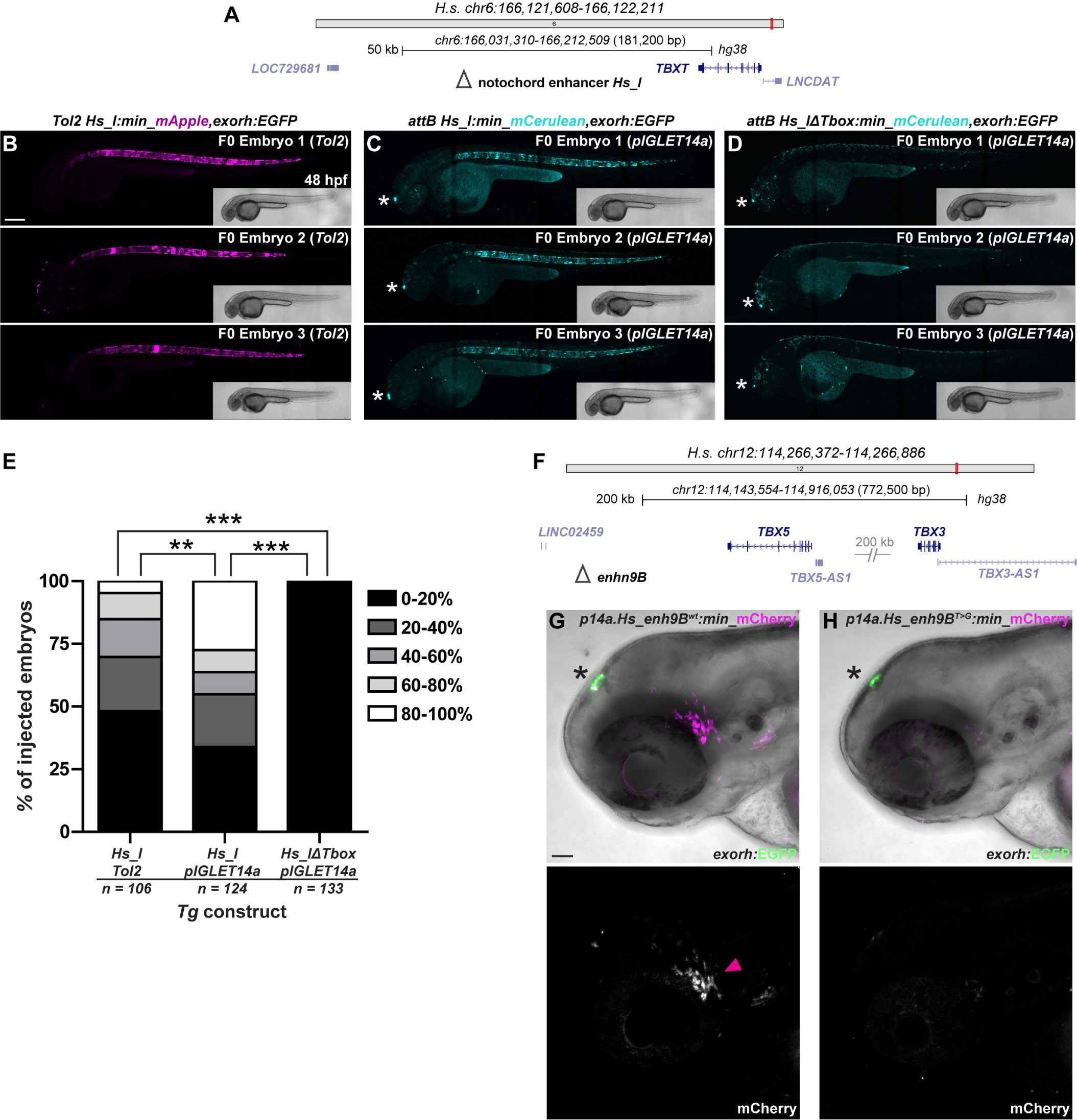
*pIGLET-*based transgenesis facilitates consistent, reproducible enhancer variant testing in Zebrafish. (**A-E**) F0 injection-based testing of notochord enhancer *Hs_I* using *Tol2-* and *pIGLET-*based transgenesis. Schematic of *TBXT* locus with *Hs_I* annotated (**A**). Confocal imaging for mApple (**B**, magenta fluorescence) or mCerulean (**C, D**, cyan fluorescence) with brightfield inserts, anterior to the left. Both *Tol2-* and *pIGLET14a-*based F0-injections of *Hs_I:min-mApple/mCerulean, exorh,EGFP* result in embryos showing reporter expression with 80-100% fluorescent notochord coverage (**B, C**). Removal of the putative Tbox motif from *Hs_I* results in a complete loss of notochord reporter activity **(D).** Injected zebrafish are sorted for the *exorh:EGFP* transgenesis marker (white asterisk) to confirm successful injection (**C, D**). *pIGLET14a*-based F0-injections of *Hs_I:min-mCerulean, exorh:EGFP,* show significantly more embryos with 80-100% fluorescent notochord coverage compared to *Tol2-*based injections (*Hs_I:min-mApple, exorh:EGFP*) and the Tbox motif deletion variant (*Hs_IΔTbox:min-mCerulean, exorh:EGFP*) (**E**). (**F-H**) Comparison of stable reporter lines driven by the human *TBX5/TBX3*-associated enhancer *9B* (*Hs_enh9B*, *wt* versus *T>G* variant). Schematic of *TBX5/3* locus with *enh9B* annotated (**F**). *Hs_enh9B^wt^* drives reporter expression in the ventral lateral line ganglia (magenta arrowhead) (**G**). Compared to *wt*, the disease-associated variant *Hs_enh9B^T>G^* exhibits nearly a complete loss of *mCherry* reporter expression (**H**) when both transgenes are integrated in the *pIGLET14a* locus (**G, H**). Note the presence of the *exorh:EGFP* transgenesis marker for screening purposes (black asterisk) (**G, H**). Scale bars: B, C, D = 200 μm, G, H = 50 μm.

We next sought to validate *pIGLET*-based enhancer testing with a human disease-associated, non-coding variant. Enhancer (*enh9B*) within the human *TBX5/TBX3* gene cluster has previously been described to harbor a SNP (*T>G*) that associates with congenital heart defects (**Fig. 4F**)^55^. In stable transgenic reporter mice, *enh9* drives reporter expression in the cardiac ventricles and more broadly throughout the embryo in regions associated with both *TBX5* and *TBX3* activity, while zebrafish reporter expression was only reported for injection-based F0 assays^55^. Stable mouse reporter transgenes for the SNP (*T>G*) showed greatly reduced or lost cardiac reporter activity, with uncharacterized impact on other expression domains^55^. After generating *attB*-based transgenesis vectors for *Hs_enh9B^wt^* from wild type reference and *Hs_enh9B^T>G^* from variant sequences for zebrafish transgenesis using *pIGLET*, we isolated stable targeted integrations for each in *pIGLET14a* (**Fig. 4G,H**). Consistent with *TBX3* expression in diverse ganglion populations in zebrafish and mice^56–59^, we observed prominent reporter activity of *Hs_enh9B^wt^:min-mCherry* in ventral anterior lateral line ganglia (**Fig. 4G**). In contrast, integrated at the same locus, the SNP version *Hs_enh9B^T>G^:min-mCherry* showed loss of activity in these ganglia (**Fig. 4H**). In either line, we did not observe any cardiac reporter expression, arguing against a conserved activity of this enhancer in the heart. Together, these results show the utility of *pIGLET*-based transgenesis in zebrafish to qualitatively assess regulatory element activity of variants associated with human disease.

## DISCUSSION

phiC31 integrase-mediated transgenesis into well-established landing sites has been transformative for work in *Drosophila* and mice^18–21^. Our proof-of-concept experiments here establish the *attP* landing sites *pIGLET14a* and *pIGLET24b* for highly efficient, reproducible, and quality-controllable transgenesis using phiC31 integrase in zebrafish. Our protocols outlined here pave the way to expand the repertoire of validated landing sites available to the model and enable critical transgene-based techniques.

Transgenesis into established *attP* landing sites in *Drosophila* and mouse hinged on identifying suitable sites by screening and validation of random integrations^18–21^. Our data here indicates that *attP*-based landing sites with favorable properties can be generated in zebrafish by re-purposing well-validated transgenes that have remained functional for numerous generations. Our approach is particularly suitable to convert existing *Tol2*-based transgenes with matching guideRNAs (**Fig. S1**). The original *Tol2*-based *ubi:Switch* integration dates back to 2009^17,29^ and *hsp70l:Switch* was isolated in 2016^17,28^, with *ubi:Switch* in use with numerous zebrafish labs since distribution started in 2010. CRISPR-Cas9-mediated conversion of *Tol2*-based transgenes to *attP* landing sites via a two-step process (or one-step if the native locus is amenable) is applicable to other well-established transgenes available in the field, and potentially any other model system.

Our demonstration of predictable, standardized, highly efficient transgene integration and suitability to generate previously challenging reporters such as *loxP* Switches establish *pIGLET14a* and *pIGLET24b* as widely applicable reagents for zebrafish transgenesis. Our provided protocols and recommended quality control steps use standard reagents and workflows established for general zebrafish transgenesis, supporting simple implementation (see also **Methods** and **Supplementary Methods**). Complementing existing random transgenesis performed with prior methods in the field (Tol2, I-SceI), our first *pIGLET* landing sites enable streamlined transgenic strain generation with reduced screening and animal numbers. Targeted, reproducible transgene integration into validated sites proposes the use of *pIGLET* for large-scale transgene tests with quantitative and qualitative readouts in zebrafish applicable to disease variant testing, mechanistic investigation of gene-regulatory elements, and more.

Being homozygous viable, our first two landing sites enable combinatorial use of maximally four individual transgenes. Future efforts to establish additional *pIGLET* landing site lines will greatly expand the possibilities for combinatorial transgene experiments and choice of tailored landing sites for specific transgene applications^22,26,27^. Strategic generation and validation of additional *pIGLET* landing sites has the potential to enable methods involving controlled mitotic chromosome arm recombination such as the Mosaic Analysis with a Repressible Cell Marker (MARCM)^60^ or Mosaic Analysis with Double Markers (MADM)^61^ systems applied in Drosophila and mice, respectively, for next-level cell labeling and conditional loss-of-function experiments. Importantly, given access to the same *pIGLET* lines, transgenic strains can be reproduced in different labs by mere sharing of the original transgenesis vector, which has the potential to drastically expand opportunities for worldwide reagent sharing in the field.

## ACKNOWLEDGEMENTS

We thank Christine Archer, Ainsley Gilbard, and Olivia Gomez for zebrafish husbandry at CU Anschutz; all past and present members of the Mosimann lab for constructive input and support during the pursuit of this long-term project; Piglet the cat for acronym inspiration; and Dr. Joaquín Navajas Acedo, Mireia Codina Tobias, and Dr. Alexander Schier for critical input and field testing.

## AUTHOR CONTRIBUTIONS

RLL and CM wrote the manuscript. RLL, HHW, CLK, RLS, AB, and CM performed plasmid cloning. RLL created the *pIGLET14a* and *pIGLET24b* landing sites. RLL created the *p14a/p24b.drl:EGFP, p14a/p24b.lmo2:EGFP, cryaa:Venus, p14a/p24b.elavl3:mCherry,exorh:EGFP, p14a/p24b.drl:creERT2,cryaa:Venus* and *p14a/p24b.hsp70l:Switch2,cryaa:Venus* transgenic lines. HHW created the *p14a*.*Hs_enh9B:min-mCherry,exorh:EGFP* transgenic lines, and RLL and CLK performed the *Hs_I:min-mCerulean;exorh:EGFP(pIGLET14a), Hs_I:min-mApple,exorh:EGFP(Tol2)* F0 analysis. SN performed preliminary testing of sgRNA combinations and proof-of-concept for Tol2 transgene remobilization. RLL and CM conceptualized the project and CM and AB provided funding.

## METHODS

### Landing site creation (2-step)

**\* Step 1: *ubi:Switch and hsp70l:Switch* transgene deletion:** sgRNAs targeting *Tol2* arms (*Tol2 ccA, Tol2 ccB*) were co-injected with a single-strand oligonucleotide ssODN containing the 84 bp *attP1* sequence and symmetric homology arms (48 bp) into *ubi:Switch* or *hsp70l:Switch* homozygous embryos (Thermo Scientific, PAGE-purified, sequence details below). Injected F0 zebrafish were raised and out-crossed to identify transgene-negative progeny, indicative of full or partial transgene deletion, or *attP1* insertion. Transgene-negative progeny was PCR-assayed (Primers: *oRL095 Tol2 remnant Fw*, *oRL096 Tol2 remnant Rev*) and sequenced to test for transgene deletion/*attP1* insertion, with the absence of any PCR product suggesting partial transgene retention. No injected zebrafish contained the *attP1* insertion. 1/20 injected zebrafish for both the *ubi:Switch* and *hsp70l:Switch locus*, transmitted F1 progeny with perfect or near perfect (single SNP) transgene deletion between the *Tol2* sgRNA target sites. Single F1 transgene deletion zebrafish were then out-crossed and heterozygous F2 zebrafish were in-crossed for the next step.

**\* Step 2: *attP1* landing site insertion:** A single sgRNA targeting the *Tol2* arm remnants (*Tol2 ccC*) was co-injected with a ssODN (2.0, sequence details below) containing the 84 bp *attP1* sequence and asymmetric homology arms (37 bp & 59 bp) into F2 heterozygous in-crossed *ubi:Switch* or *hsp70l:Switch* transgene deletion animals. F0 zebrafish were raised, fin clipped, PCR-assayed (Primers: *oRL095 Tol2 remnant Fw*, *oRL096 Tol2 remnant Rev*) and sequenced to test for evidence of *attP1* insertion. Importantly, F0 zebrafish that displayed *attP1* sequence insertion in the somatic tissue (fin clip) invariantly transmitted the same alleles in the germline, indicating homology directed repair occurred early during development. From zebrafish that contained the *Tol2* arm remnant target site, 1/18 showed *attP1* insertion at the *ubi:Switch* locus, and 1/35 showed *attP1* insertion at the *hsp70l:Switch* locus. Single F1 heterozygous zebrafish (now termed *pIGLET14a* and *pIGLET24b*) were out-crossed, and heterozygous F2 zebrafish were in-crossed to obtain F3 homozygous *pIGLET14a* and *pIGLET24b* zebrafish.

### RNP complex preparation

Injection mixes for both step 1 and step 2 were made as per our previous work^30^ using the following final concentrations: 0.8 μg/μL Cas9 (Alt-R™ S.p. Cas9 Nuclease V3, 100 µg – IDT #1081058), ssODN 60 ng/μL, 0.05% phenol red, 300mM KCl, make up total volume with sgRNA.

### sgRNA target sites

*Tol2 ccA: 5’-GGGCATCAGCGCAATTCAAT-3’*

*Tol2 ccB: 5’-GTGTATTAGTCTTGATAGAG-3’*

*Tol2 ccC: 5’-AGTGCTGAAAAGCCTCTCAC-3’*

### ssODN sequences

*attP1* (bold/underlined) ssODN: *5’- TTTGGAGATCACTTCATTCTATTTTCCCTTGCTATTACCAAACCAATT**AGAAGCGGTTTTCGGGAGTAG TGCCCCAACTGGGGTAACCTTTGAGTTCTCTCAGTTGGGGGCGTAGGGTCGCCGACATGACAC**TATC AAGACTAATACACCTCTTCCCGCATCGGCTGCCTGTGAGAGGCT-3’*

*attP1* (bold/underlined) ssODN 2.0: *5’- ATCAAGACTAATACACCTCTTCCCGCATCGGCTGCCT**AGAAGCGGTTTTCGGGAGTAGTGCCCCAAC TGGGGTAACCTTTGAGTTCTCTCAGTTGGGGGCGTAGGGTCGCCGACATGACAC**GTGAGAGGCTTTT CAGCACTGCAGGATTGCTTTTCAGCCCCAAAAGAGCTAGGCTTGAC-3’*

### Primer sequences

*oRL095 Tol2 remnant Fw*: *5’-ATTGCCAGAGGTGTAAAGTA-3’*

*oRL096 Tol2 remnant Rev*: *5’-CCAGTACACGCTACTCAAA-3‘*

### pIGLET14a, pIGLET24b genotyping

Unrecombined, native *attP1* alleles: *oRL095 Tol2 remnant Fw* + *oRL096 Tol2 remnant Rev* (expected size ∼500-520 bp, depending on *pIGLET14a or pIGLET24b*)

**→ PCR program**
30 cycles
Tm = 54 °C
Extension = 30 sec

*pIGLET14a*^+/+^ or ^-/+^ genotyping: *oRL168 ubi:Switch locus Fw* + *oRL169 ubi:Switch locus Rev* (Expected size 1448bp for presence of landing site, 631bp for absence of landing site)

**→ PCR program**
30 cycles
Tm = 65 °C
Extension = 1 min

*oRL168 ubi:Switch locus Fw: 5’-CATGATCGAAAACGAGCAATGTC-3’*

*oRL169 ubi:Switch locus Rev: 5’-GACTGCGTCACTTTGACAACCCTG-3’*

*pIGLET24b*^+/+^ or ^-/+^ genotyping: *oRL074 hsp70l:Switch locus Fw* + *oRL073 hsp70l:Switch locus Rev* (Expected size 1421 bp for presence of landing site, 627 bp for absence of landing site)

**→ PCR program**
30 cycles
Tm = 57 °C
Extension = 1 min

*oRL074 hsp70l:Switch locus Fw*: *5’-GCATGACACGGCTAACCAAC-3’*

*oRL073 hsp70l:Switch locus Rev*: *5’-TATCAGCACACACCTTTATCGC -3’*

### Transgenic line creation (additional details contained in Supplementary Methods)

Single cell *pIGLET14a* and *pIGLET24b* embryos were co-injected with 25 ng/μL of expression plasmid (containing phiC31 recognized-compatible *attB* recombination cassette) and 25 ng/μL of phiC31 *integrase* mRNA. 10-20 F0 embryos/construct were sacrificed to check for targeted integration as described in the **Supplementary Methods** (printable for bench use). ∼25-50 % of transmitted F1 embryos from the first transgene-positive clutch were genotyped to check for correct integration. Only clutches that showed 100% targeted integration were raised. F1 adults were genotyped by fin clip to check for targeted integration and out-crossed to check for expected Mendelian transmission ratios in the next generation (50% transgene-positive), indicative of single copy transgene insertions.

For detailed integration methods, including sample gels and quality control suggestions, and steps to create DNA sequence map of plasmid integrations into *pIGLET14a* and *pIGLET24b* loci, please see **Supplementary Methods**. Of note, if *pRL092 pattB_cryaa:Venus_MCS* or *pRL093 pattB_exorh:EGFP_MCS* were used for expression plasmid construction, a customized *Fw* primer will be needed in place of primer *attB2 Fw* to sequence-confirm the cloned 3’ integration boundary.

### phiC31 integrase mRNA

*pCDNA3.1 phiC31 integrase* (Addgene #68310)^20,22^ plasmid was linearized with BamHI and transcribed using the T7 mMessage mMachine kit (ThermoFisher AM1344). Following IVT, mRNA was treated with Turbo DNase cleaned using LiCl precipitation methods as described in the T7 mMessage mMachine kit protocol. Briefly, following Turbo DNAse treatment, 30 μL of nuclease-free water and 30 μL of LiCl was added and the reaction was precipitated overnight at -20 °C. Following a 15 min max speed centrifuge, 750 μL of cold 70% EtOH was added, and the reaction was spun again for 10 min at max speed. After careful removal of EtOH, the pellet was air dried and resuspended in 20 μL of nuclease-free water. Clean preps of phiC31 *integrase* mRNA are well-tolerated by zebrafish embryos; if excessive mortality is observed, IVT of a new mRNA batch and careful cleanup is recommended.

### 4-OHT inductions, heat-shock treatments

Activity of CreERT2 was induced with 10 μM final concentration of (Z)-4-Hydroxytamoxifen (Sigma Aldrich, St. Louis, Missouri, H7904, abbreviated as 4-OHT) in E3^62^. 4-OHT stock is stored at −20 °C in the dark as 10 mM single-use aliquots dissolved in DMSO and used within 2 months of dissolving. Prior to administration, the 4-OHT aliquots were incubated at 65 °C for 10 minutes and vortexed^62^. For shield stage treatment, 4-OHT was administered overnight and then replaced with N-Phenylthiourea (Sigma Aldrich, P7629, abbreviated as PTU) at a final concentration of 150 μM in E3 embryo medium each morning to inhibit melanogenesis.

To trigger fluorophore expression, *hsp70l:Switch* and *p14a/p24b.hsp70l:Switch2* embryos were heat-shocked for 1 hour at 37 °C in a dedicated water bath. Prior to heat shock, embryos were dechorionated and transferred to a glass vial with E3 medium. Embryos were imaged 3-4 hours (confocal microscope) later.

### Genomic DNA isolation

To isolate genomic DNA, embryos/fin clips are sorted into 0.2 μL PCR tubes containing 50 μL of 50mM NaOH solution. Using a thermocycler, tubes are then heated for 20 min at 95 °C and then cooled to 12°C. Afterwards, 5 μL of 1.0 M Tris buffer (pH 8.0) is added and genomic preps are stored at 4 °C. 1-2 μL are used for a typical PCR reaction.

### Plasmid construction

All plasmids were created with the Multisite Gateway system with LR Clonase II Plus (Cat#12538120; Life Technologies) according to the manufacturer’s instructions and original Tol2 Kit information^2,31^, assemblies described below and vector ratios calculated using the Multisite Gateway Excel Spreadsheet^63^:

***pCM318 drl:EGFP***: *pCM293 pENTR5’_*6.3kb *drl* regulatory region^32^, *#383 pENTR/D_EGFP*^2^, *#302 p3’_SV40 polyA*^2^, *pCM268 pDESTattB*^22^ (Addgene #68313).

***pCM328 lmo2:EGFP,cryaa:Venus***: *pENTR5’_lmo2* upstream region^36^, *#383 pENTR/D_EGFP*^2^, #302 *p3’_SV40 polyA*^2^, *pCM327 pDESTattB_cryaa:Venus*^22^ (Addgene #68341).

***pRL058 elavl3:mCherry,exorh:EGFP:*** *pENTR5’_8.7 kb elavl3(HuC)*^38^, #456 *pENTR/D_*mCherry^2^, *pGD003 p3’_ubbpA*^31^, *pRL056 pDESTattB_exorh:EGFP*.

***pRL045 drl:creERT2, cryaa:Venus:*** *pCM293 pENTR5’_*6.3kb *drl* regulatory region^32^, *pENTR/D_creERT2* (Addgene #27321)^29^, *pGD003 p3’_ubbpA*^31^, *pRL055 pDESTattB_cryaa:Venus*.

***pRL069 hsp70l:Switch2,cryaa:Venus:*** *pDH083 pENTR5’*_*hsp70l:loxP-Stop-loxP*^64^, *#763 pENTR/D_mApple*^31^, *pGD003 p3’_ubbpA*^31^, *pRL055 pDESTattB_cryaa:Venus*

***pRL050 Hs_enh9B^wt^:min-mCherry,exorh:EGFP*:** *pRL049 pENTR5’_Hs_enh9B^wt^*, *pCK006 pENTR/D_min-mCherry*^31^*, pGD003 p3’_ubbpA*^31^*, pRL056 pDESTattB_exorh:EGFP*

***pRL052 Hs_enh9B^T>G^:min-mCherry, exorh:EGFP*:** *pRL051 pENTR5’_Hs_enh9^T>G^*, *pCK006 pENTR/D_min-mCherry*^31^*, pGD003 p3’_ubbpA*^31^*, pRL056 pDESTattB_exorh:EGFP*

***pCK086 Hs_I:min-mCerulean, exorh:EGFP:*** *pENTR5’_Hs_I*^54^*, pSN001 pENTR/D_min-mCerulean*^31^, #302 *p3’_SV40 polyA*^2^*, pRL056 pDESTattB_exorh:EGFP*

***pCK085 Hs_IdelTbox:min-mCerulean, exorh:EGFP:*** *pCK075 pENTR5’_Hs_IdelTbox*^54^*, pSN001 pENTR/D_min-mCerulean*^31^, #302 *p3’_SV40 polyA*^2^*, pRL056 pDESTattB_exorh:EGFP*

### pENTR5’ pRL049 Hs_enh9B^wt^ *and* pENTR5’ pRL051 Hs_enh9B^T>G^ *cloning*

*pRL049 Hs_enh9B^wt^* was amplified with the following primers, using human genomic DNA as template, and then cloned into the pENTR5’-TOPO vector (Thermo Fisher #K59120):

*oRL163 Hs_enh9B Fw*: *5’-AGTAGGGGGATGCTAATTTCATAGC-3’*

*oRL164 Hs_enh9B Rev*: *5’-TCTCAAGTCCTCTTGCGGTTTTGA-3’*

*pRL051 Hs_enh9^T>G^* was cloned using *pRL049* as a template for overlapping PCR to introduce the *T>G* mutation at base pair position 339 using the following primers:

*oRL165 Hs_enh9B overlap Rev: 5’-CCCAACAGTTTCCAAGCATATTGAATATATTGTGGTCAG-3’*

*oRL166 Hs_enh9B overlap Fw: 5’-CTGACCACAATATATTCCATATTCTTGGAAACTGTTGGG-3’*

The products of *oRL163 Fw* + *oRL65 Rev* and *oRL166 Fw* + *oRL164 Rev* were individually column-purified and 1 µl of each was combined as template for a final PCR reaction using *oRL163 Fw* + *oRL164 Rev*. The final product was then cloned into the *pENTR5’-TOPO* vector (Thermo Fisher #K59120).

### Imaging

PTU-treated (150μM) embryos were anesthetized at 10 ss - 2 dpf with 0.016 % Tricaine-S (MS-222, Pentair Aquatic Ecosystems, Apopka, Florida, NC0342409) in E3 embryo medium. Standard fluorescence imaging was performed on a Leica M205FA with a DFC450 C camera. Laser scanning confocal microscopy was performed on a Zeiss LSM880 following embedding in E3 with 1 % low-melting-point agarose (Sigma Aldrich, A9045) on glass bottom culture dishes (Greiner Bio-One, Kremsmunster, Austria, 627861). Images were collected with a ×10/0.8 and x20/0.8 air-objective lens with all channels captured sequentially with maximum speed in bidirectional mode, with the range of detection adjusted to avoid overlap between channels. Maximum projections of acquired Z-stacks were made using ImageJ/Fiji and cropped and rotated using Adobe Photoshop.

### Statistics

For Figure 3D, each category of notochord fluorescent coverage was scored between 1 and 5 (0-20% = 1, 20-40% = 2, 40-60% = 3, 60-80% = 4, 80-100% = 5) and the sum/average of the fluorescent coverage scores per group were calculated. The average fluorescent coverage score of each group was then compared to each other using Kruskal-Wallis test with Dunn’s correction for multiple tests. The proportion of embryos presenting each fluorescent coverage score per group is included. Adjusted *P* values after multiple tests correction are reported and significance was set at *P* < 0.0001. GraphPad Prism 9.0.2 was used to perform statistical tests and generate graphs.

---

**Figure S1.**
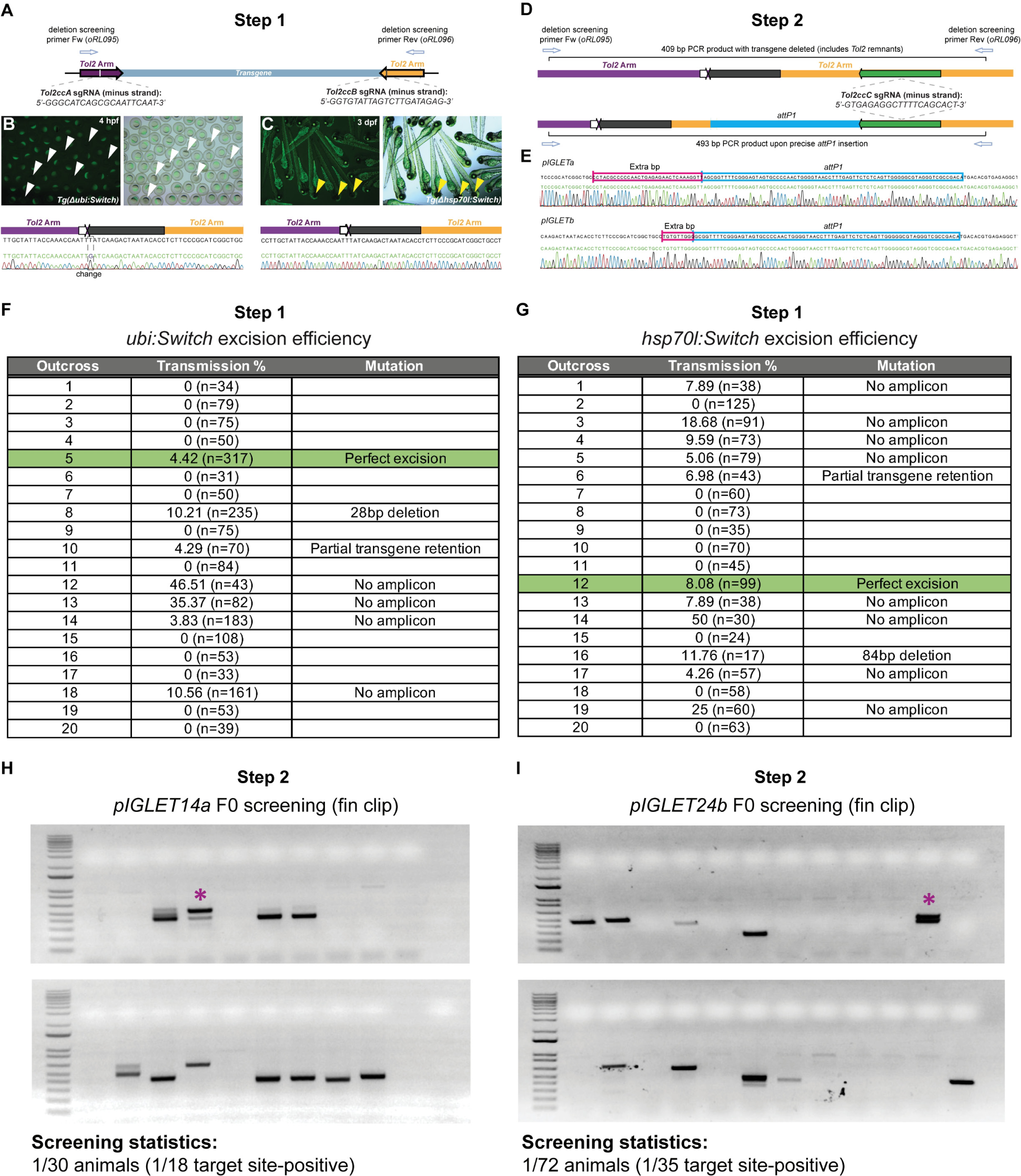
Schematic of *pIGLET14a* and *pIGLET24b* landing site allele design and allele screening statistics. **Step 1:** (**A**) *ubi:Switch* and *hsp70l:Switch* transgenes were deleted using sgRNAs targeting the *Tol2* arms. (**B, C**) Step 1: Injected F0 homozygous *ubi:Switch* or *hsp70l:Switch* zebrafish were out-crossed and F1 progeny were screened for loss transgene expression indicative of full or partial transgene deletion. Loss of ubiquitous *EGFP* expression (white arrowheads) or loss of *cryaa:Venus* expression (yellow arrowheads) indicated deletion of *ubi:Switch* and *hsp70l:Switch* transgenes, respectively, subsequently confirmed by PCR and sequencing (**A, B**). Note partial *Tol2* arm remnants following transgene deletion (**A, B**). (**D, E**) Step 2: Following transgene deletion, a single *Tol2* remnant-specific sgRNA was co-injected with a ssODN containing the *attP1* landing site and homology arms to create the *pIGLET14a* and *pIGLET24b* alleles. (**F, G**) Step 1: One out of 20 F0-injected zebrafish transmitted F1 embryos with complete transgene excision for both the *ubi:Switch* and *hsp70l:Switch* transgenic lines. Note F0-injected fish transmitted imperfect deletion F1 embryos (*Δ28bp*, *Δ84bp*), minor transgene retention (obtained amplicon), and major transgene retention (no amplicon) (**F, G**). (**H, I**) Step 2: Following the second round of sgRNA + *attP1* ssODN injection, F0-injected zebrafish were fin clipped to detect somatic *attP1* insertion events, which ultimately were transmitted in the germline. F0 fin clips showing insertion events that led to *pIGLET14a* and *pIGLET24b* alleles (magenta asterisk), and screening statistics (**H, I**).

**Figure S2.**
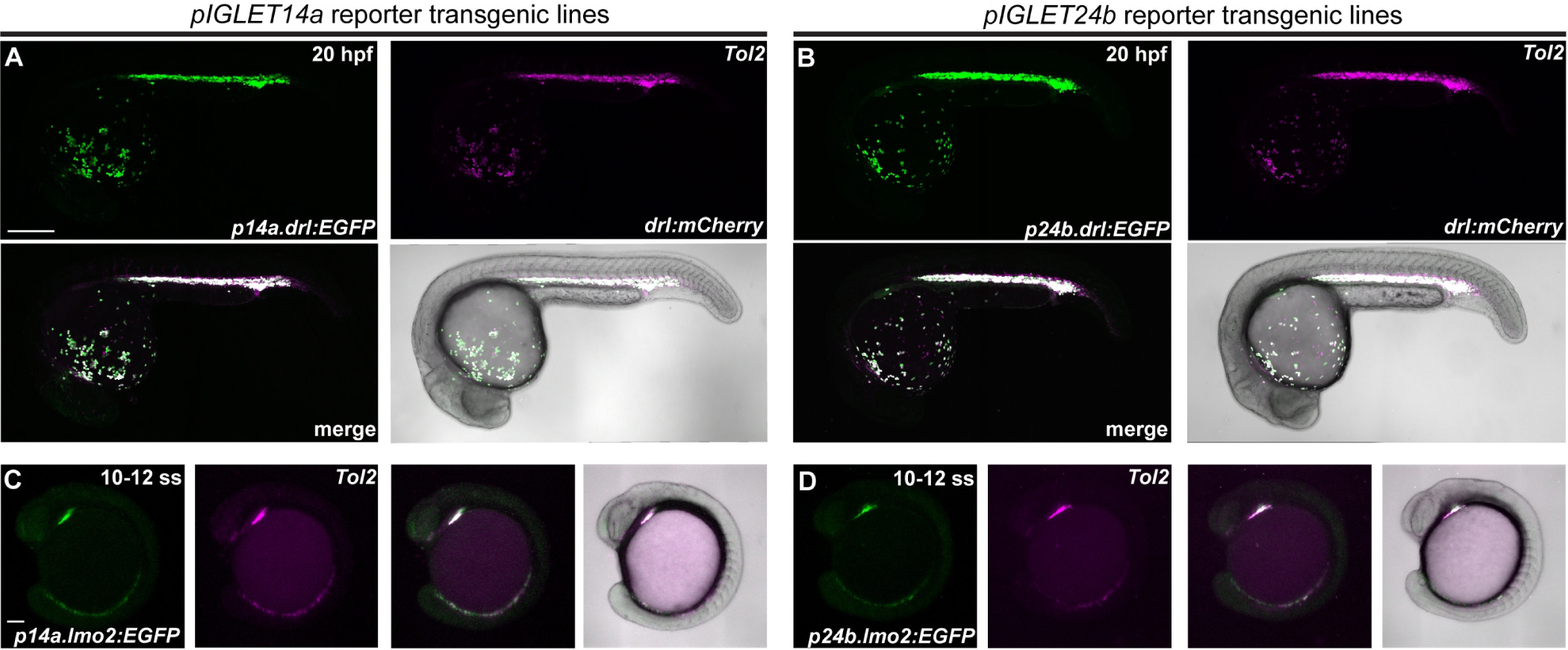
Extended reporter transgene testing in *pIGLET14a* and *pIGLET24b* landing sites. (**A-D**) Continued validation of *pIGLET14a* and *pIGLET24b* transgenics for *drl:EGFP* and *lmo2:EGFP* from Figure 1. Confocal images, anterior to the left. When crossed to d*rl:mCherry(Tol2*), *p14a*.*drl:EGFP* and *p24b.drl:EGFP* show faithful, overlapping reporter activity in hematopoietic lineages at 20 hpf, consistent with previously published *Tol2*-based *drl:EGFP* transgenic lines (**A, B**). When crossed to *lmo2:Switch* (*loxP-dsRED2-loxP_EGFP*) as reference, *p14a.lmo2:EGFP* and *p24b.lmo2:EGFP* show faithful, overlapping reporter activity in the endothelial progenitor populations at 10-12 ss (**C, D**). Scale bars: A, B = 250; μm, C, D = 100 μm.

**Figure S3.**
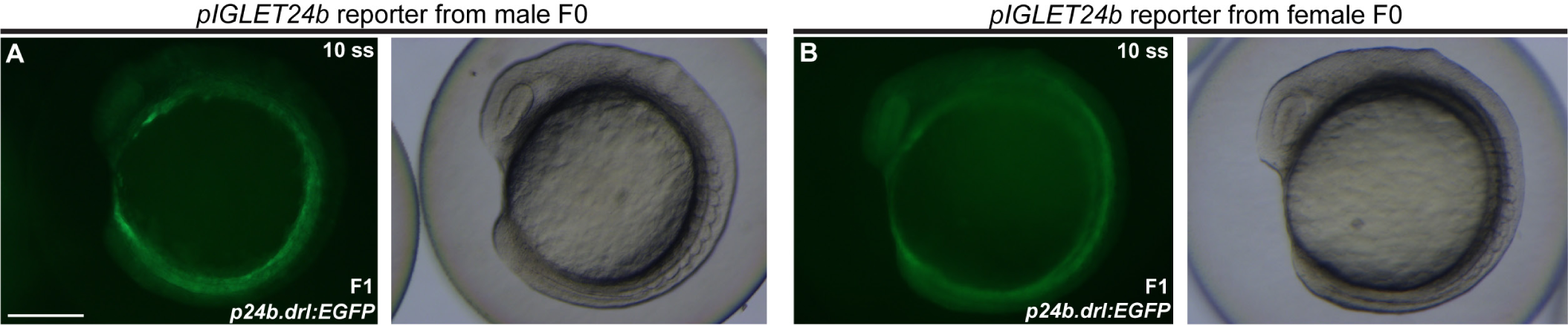
Extended *p24b.drl:EGFP* transgene testing in *pIGLET24b* landing sites. (**A, B**) At the *pIGLET24b* locus, *drl:EGFP* display activity maternal expression, as observed by ubiquitous *EGFP* in *p24b.drl:EGFP* when embryos are derived from female transgene transmission; note the difference in expression of *EGFP* in *p24b.drl:EGFP* F1s from male and female parents. Dissecting scope images, anterior to the top, ventral to the left. Scale bars: A, B = 215 μm.

**Figure S4.**
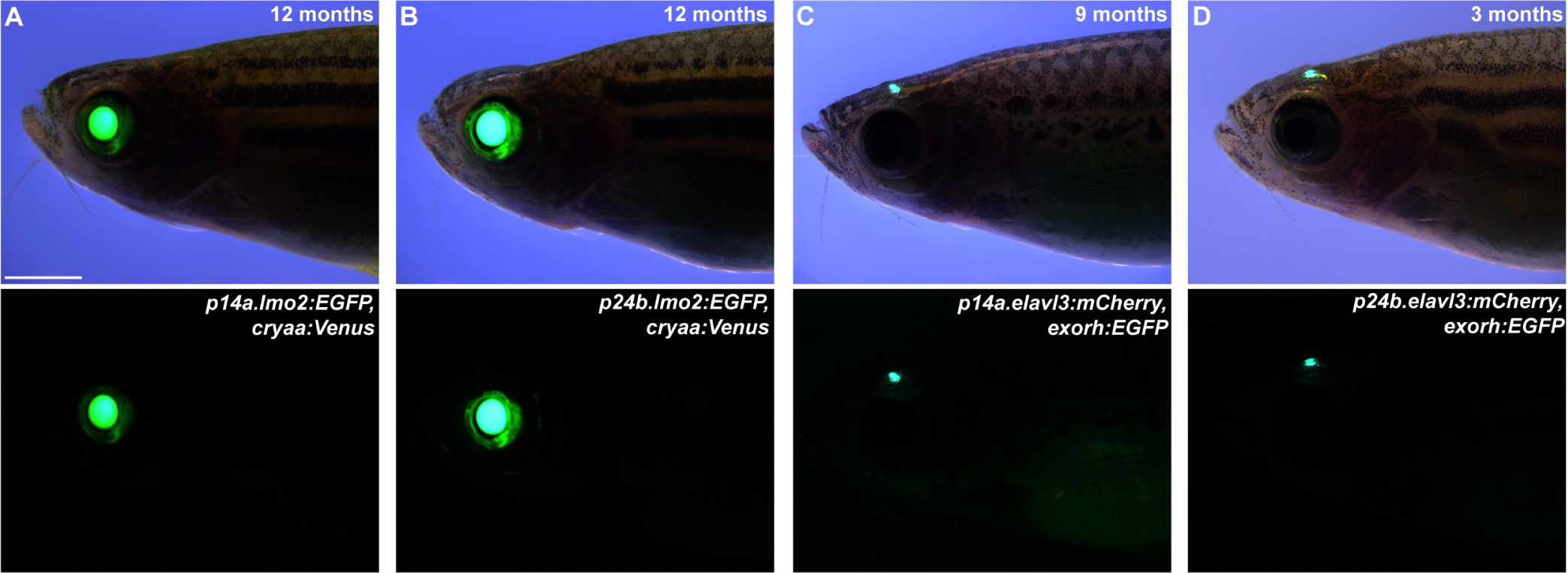
Extended adult transgene testing in *pIGLET14a* and *pIGLET24b* landing sites. (**A-D**) Persistent activity of transgenesis markers through adulthood for transgenes integrated in *pIGLET14a* and *pIGLET24b*. Adult zebrafish imaged on dissecting scope, anterior to the left. Expression of transgenesis markers *cryaa:Venus* and *exorh:EGFP* persist in adult *p14a.lmo2:EGFP, cryaa:Venus* and *p24b.lmo2:EGFP, cryaa:Venus* (**A, B**), and *p14a.elavl3:mCherry, exorh:EGFP* and *p24b.elavl3:mCherry, exorh:EGFP (***C, D**) transgenic zebrafish, respectively. Scale bars: A, B = 2.1 mm; C = 2.4 mm; D = 1.6 mm.

**Figure S5.**
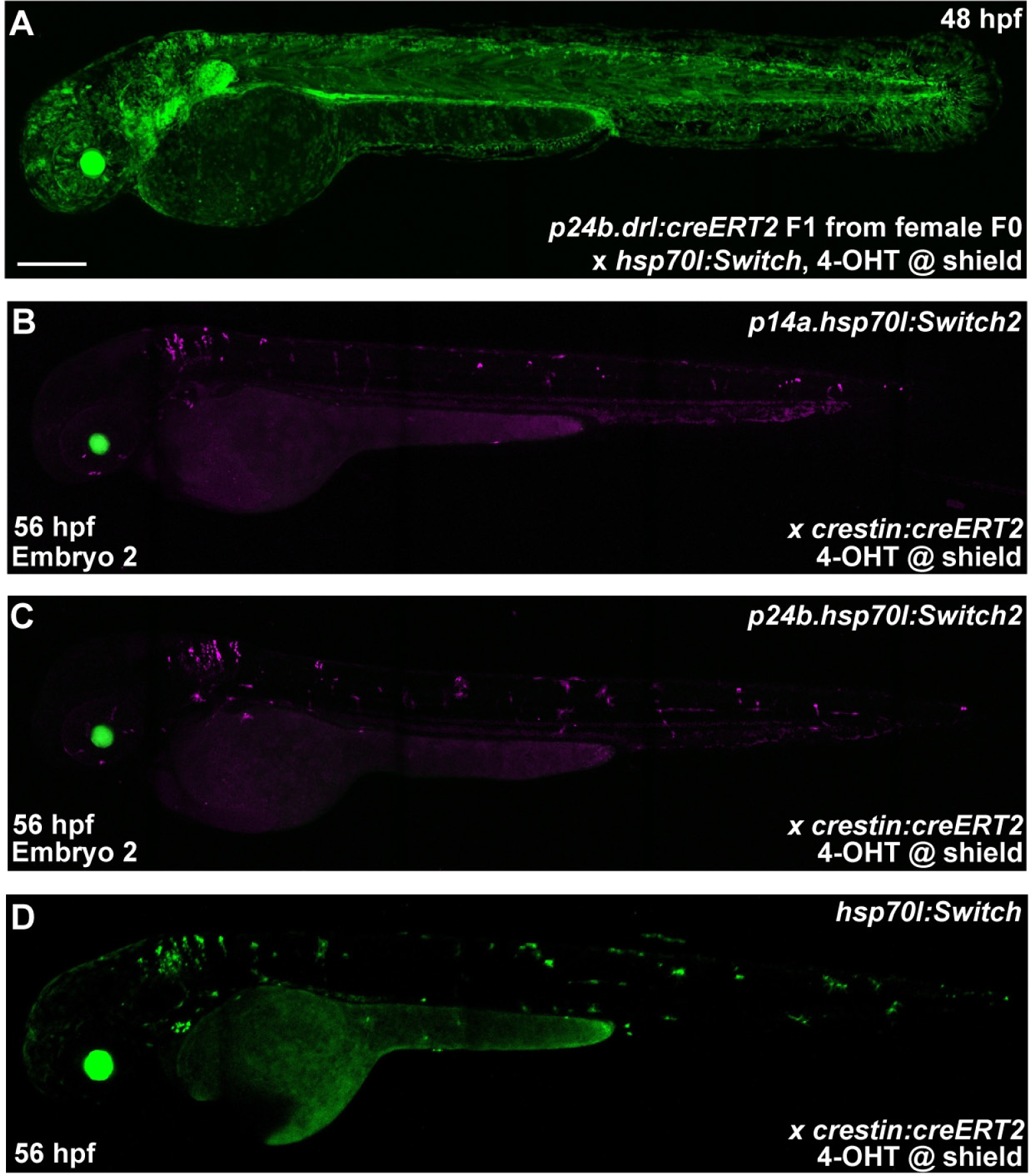
Extended CreERT2-driver testing in *pIGLET14a* and *pIGLET24b* landing sites. (**A-D**) Additional datapoints on Cre/lox-based transgenes. Confocal imaging, anterior to the left. In accordance with our *p24b*.*drl:EGFP* reporter lines, *p24b*.*drl:creERT2* embryos from a female parent show maternal CreERT2 contribution resulting in broad switching when crossed to *hsp70l:Switch* (**A**). Consistent with previously established guidelines, male CreERT2-driver zebrafish should always be used for standard Cre/*loxP*-based lineage trace experiments due to the possibility of maternal *creERT2* transcripts (**A**)^4^. When crossed to *p14a.hsp70l:Switch2* and *p24b.hsp70l:Switch2, crestin:creERT2* show recombination consistent (**B, C**) with the previously described *hsp70l:Switch* transgenic line (**D**). Scale bars: A, B, C, D = 200 μm.

**Fig. S6.**
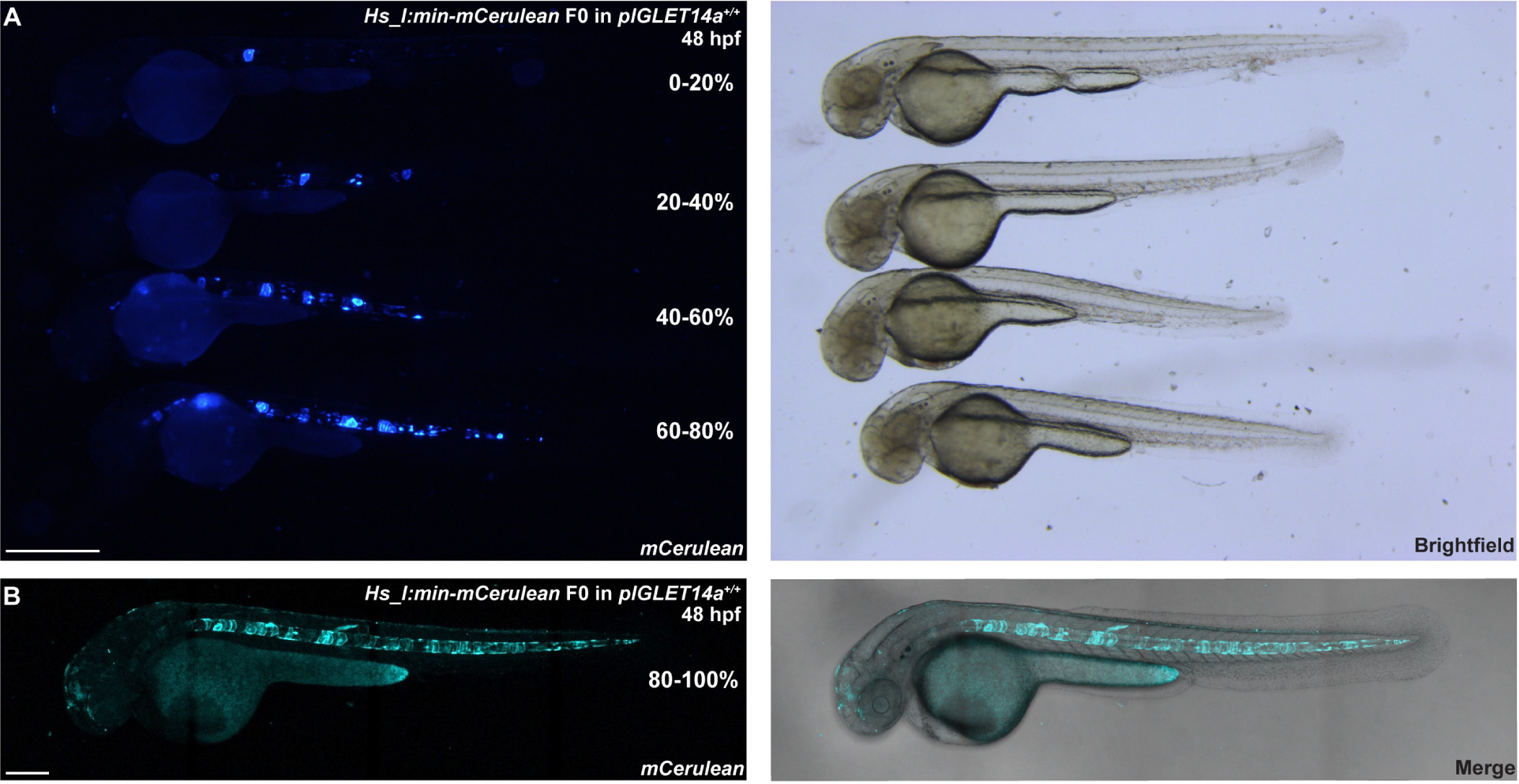
F0 notochord fluorescent coverage scoring of *Hs_I* based reporter injections. (**A-B**) Representative embryos displaying 5 subcategories of notochord fluorescent coverage: 0-20%, 20-40%, 40-60%, 60-80%, and 80-100%. Embryos shown are homozygous *pIGLET14a* injected with *Hs_I:min-mCerulean* and imaged at 48 hpf with the stereoscopic (**A**) and confocal microscope (**B**). Anterior to the left. Scale bars: A = 860 μm; B = 500 μm.

**Fig. S7.**
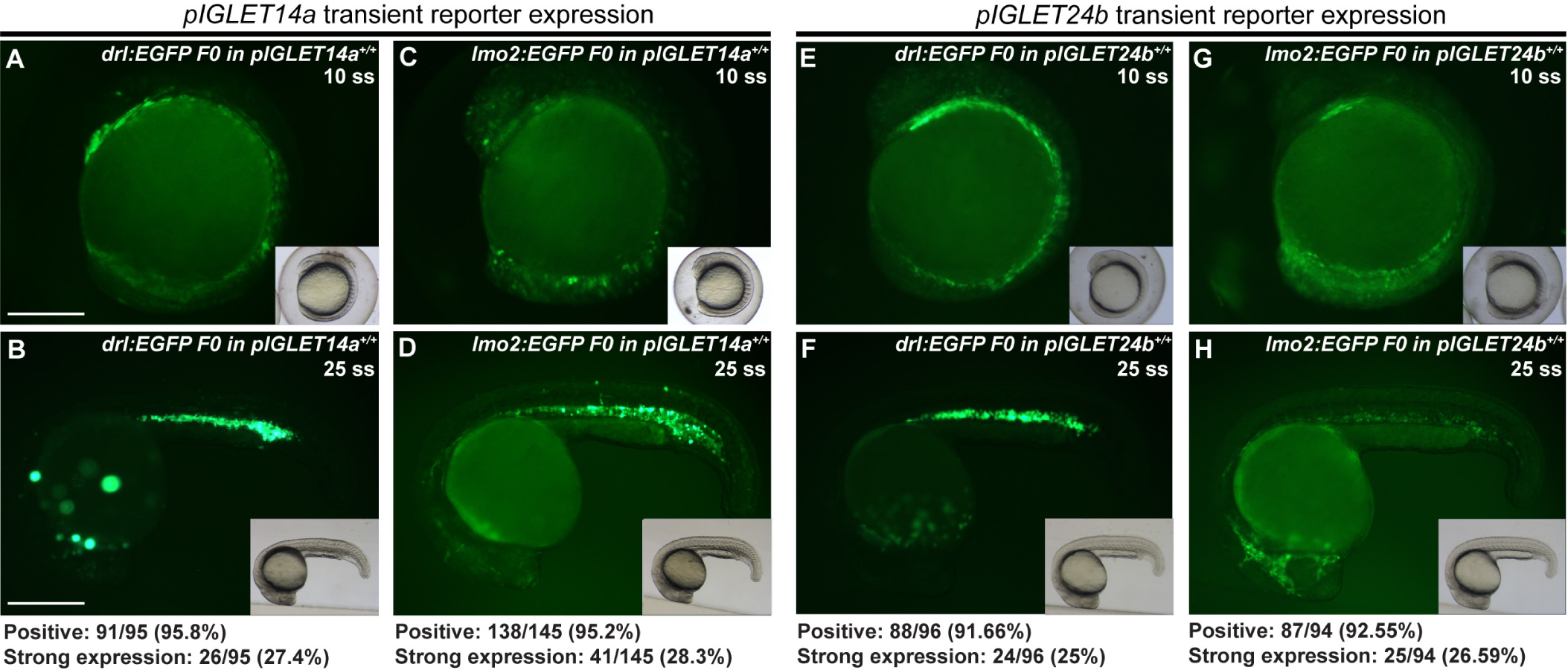
F0 *drl:EGFP* and *lmo2:EGFP* reporter injections. (**A-H**) Confocal imaging of injected embryos, anterior to the left. F0-injected homozygous *pIGLET14a* and *pIGLET24b* zebrafish show 90-95% *EGFP*-positive embryos when injected with *drl:EGFP* and *lmo2:EGFP*, with a subset (approximately 25%) showing strong *EGFP* expression approaching that of stable transgenic lines at 10 ss (**A, C, E, G**) and 25 ss (**B, D, F, H**). Scale bars: A, C, E, G = 255 μm; B, D, F, H = 310 μm.

**Table S1.** Plasmid table. Table of *pENTR5’* and expression plasmids used in this study. Description of expression plasmid assemblies is included, and specific plasmid sources are referenced in the Methods section.

| Plasmid ID | Plasmid name | Description |
| --- | --- | --- |
| **5' Entry vectors (p5E, KanR)** |  |  |
| pRL049 | *TBX5_enh9B<sup>wt</sup>* | Human regulatory element, ~88kb downstream of *TBX5* (wt variant) |
| pRL051 | *TBX5_enh9B<sup>T>G</sup>* | Human regulatory element, ~88kb downstream of *TBX5* (T>G variant) |
| **Expression Vectors (AmpR)** |  |  |
| pCM318 | *drl:EGFP* | pCM293 pENTR5' 6.3kb *drl* regulatory region, #383 pENTR/D_EGFP, #302 p3'_SV40 polyA, pCM268 pDESTattB (Addgene #68313) |
| pCM328 | *lmo2:EGFP, cryaa:Venus* | pENTR5' *lmo2* upstream region, #383 pENTR/D_EGFP, #302 p3'_SV40 polyA, pCM327 pDESTattB_ *cryaa:Venus* (Addgene #68341) |
| pRL045 | *drl:creERT2, cryaa:Venus* | pCM293 pENTR5' 6.3kb *drl* regulatory region, pENTR/D_ *creERT2* (Addgene #27321), pGD003 p3'_ *ubbpA*, pRL055 pDESTattB_ *cryaa:Venus* |
| pRL050 | *Hs_TBX5_enh9B<sup>wt</sup>:min-mCherry, exorh:EGFP* | pRL049 pENTR5' *TBX5_enh9B<sup>wt</sup>*, pCK006 pENTR/D_ *min-mCherry*, pGD003 p3'_ *ubbpA*, pRL056 pDESTattB_ *exorh:EGFP* |
| pRL052 | *Hs_TBX5_enh9B<sup>T>G</sup>:min-mCherry, exorh:EGFP* | pRL051 pENTR5' *TBX5_enh9B<sup>T>G</sup>*, pCK006 pENTR/D_ *min-mCherry*, pGD003 p3'_ *ubbpA*, pRL056 pDESTattB_ *exorh:EGFP* |
| pRL058 | *elavl3:mCherry, exorh:EGFP* | pENTR5' 8.7 kb *elavl3*(HuC), #456 pENTR/D_ *mCherry*, pGD003 p3'_ *ubbpA*, pRL056 pDESTattB_ *exorh:EGFP* |
| pRL069 | *hsp70l:Switch2, cryaa:Venus* | pDH083 pENTR5' *hsp70l:loxP-Stop-loxP*, #763 pENTR/D_ *mApple*, pGD003 p3'_ *ubbpA*, pRL055 pDESTattB_ *cryaa:Venus* |
| pCK085 | *Hs_IdelTbox:min-mCerulean, exorh:EGFP* | pCK075 pENTR5' *Hs_IdelTbox*, pSN001 pENTR/D_ *min-mCerulean*, #302 p3'_SV40 polyA, pRL056 pDESTattB_ *exorh:EGFP* |
| pCK086 | *Hs_I:min-mCerulean, exorh:EGFP* | pENTR5' *Hs_I*, pSN001 pENTR/D_ *min-mCerulean*, #302 p3'_SV40 polyA, pRL056 pDESTattB_ *exorh:EGFP* |

**Table S2.**
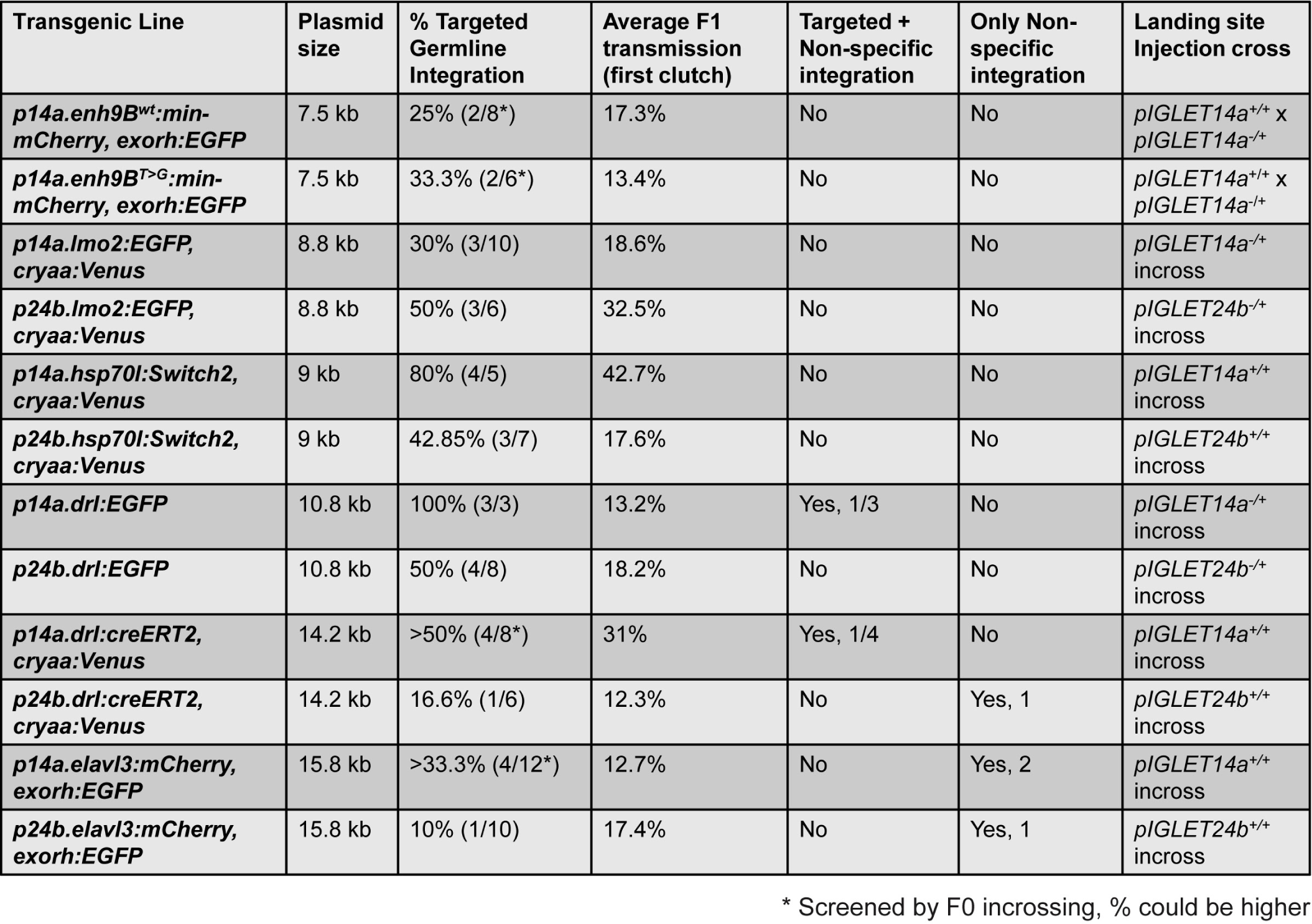
*pIGLET14a* and *pIGLET24b* transgenic line screening statistics. Full record of *pIGLET14a* and *pIGLET24b* transgenic line screening including percentage of founders obtained with targeted integrations, F1 transmission averages, instances of non-targeted transgene integration, as well as specifics of parental genotypes for plasmid injections and plasmid total size.

