## Supplementary Methods for "*pIGLET*: Safe harbor landing sites for reproducible and efficient transgenesis in zebrafish"

### Supplementary Methods: Quick reference *pIGLET* system implementation

*This document is meant as printable, bench-friendly summary of key methods associated with phiC31 integrase-mediated transgenesis into pIGLET landing sites in zebrafish, outlined in Lalonde et al.. Mosimann lab, CU Anschutz, 2023.*

#### 1. Injection

Inject 25 ng/µl plasmid + 25 ng/ul phiC31-*integrase* mRNA

- Ideal cross: *pIGLET14a* homozygous in-cross or *pIGLET24b* homozygous in-cross
- Quality control (QC): Genotype 10-20 F0 embryos to check for correct integration

**3' Integration check PCR program** (*attB1 Fw* or *attB2 Fw* + *oRL096 Tol2 remnant Rev*)

30-35 cycles (F1 vs. F0 integration check)

Tm = 54 °C

Extension = 30sec-1min (Depends on size of Gateway assembly)

*attB1 Fw: 5'-CAAGTTTGTACAAAAAAGCAG-3'*

*attB2 Fw: 5'-ACCCAGCTTTCTTGTACAAAGTGG-3'*

*oRL096 Tol2 remnant Rev: 5'-CCAGTACACGCTACTCAAA-3'*

**5' Integration check PCR program** (*oRL095 Tol2 remnant Fw* + *oRL188 5' expression plasmid Rev*)

30-35 cycles (F1 vs. F0 integration check)

Tm = 54 °C

Extension = 30 sec

*oRL095 Tol2 remnant Fw: 5'-ATTGCCAGAGGTGTAAAGTA-3'*

*oRL188 5' expression plasmid Rev: 5'-GCCTTTGAGTGAGCTGATACC-3'*

**Sample gel F0 3' Integration check (*attB2 Fw* + *oRL096 Rev*)**

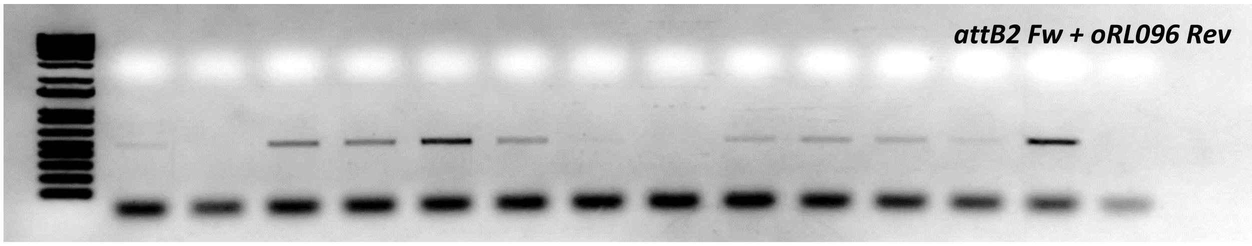


Suggestion(s):

- Increase to 35 cycles for this PCR (to increase sensitivity with mosaic integration in F0)
- Depending on type of cross, the user may want to check for presence of correct integration and presence of *attP* site (ie. no integration band expected in some embryos from +/- in-cross or outcross)
- Note, if *pRL092*, or *pRL093* were used for expression plasmid construction, a customized *Fw* primer will be needed in place of *attB1*/*attB2 Fw* to check the 3' integration boundary.

### 2. Transgenic screening

Out-cross or in-cross F0s to screen for germline transmission (All F1s will be validated anyways)

- Sort/raise by expression pattern (should all be identical)
- QC: Genotype select embryos to check for correct integration

**Sample gel F1 3' Integration check (*attB1 Fw + oRL096 Rev, attB2 Fw + oRL096 Rev*)**

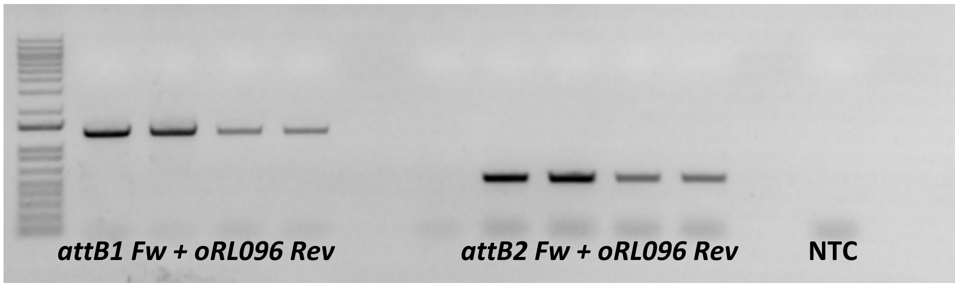


### 3. Validating alleles in adults

Fin clip individual adult F1s and genotype for correct integration

Outcross adult Fttp11s with correct integration to observe mendelian ratios (ie. check for single integration)

Suggestion(s):

- Raise F2s from F1 outcross, not in-cross (clonal expansion cleaner)
- Absolutely necessary if you screened for transgenics by in-crossing F0s, otherwise risk carrying along empty *attP* allele in transgenic line

**Integration boundary sequences (*attP*, *attB*)**

**5' boundary**

***AGCGGTTTTCGGGAGTAGTGCCCCAACTGGGGTAACCTTTGGGCTCCCCGGGCGCGTACTCCACCTCA***

***CCCATCTGGTCCATCATGATGAACGGGTCGAGGTGGCGGTAGTTGATCCCGGCGAACGCGCGGCGCA***

***CCGGGAAGCCCTCGCCCTCGAAACCGCTGGGCGCGGTGGTCACGGTGAGCACGGGACGTGCGACGG***

***CGTCGGCGGGTGCGGATACGCGGGGCAGCGTCAGCGGGTTCTCGACGGTCACGGCGGGCATGTCGAC***

**3' boundary**

***GTCGACGATGTAGGTCACGGTCTCGAAGCCGCGGTGCGGGTGCCAGGGCGTGCCCTTGAGTTCTCTCA***

***GTTGGGGGCGTAGGGTCGCCGACATGACAC***

**Steps to manually introduce plasmid into landing site locus in genomics software:**

1. Create a linear file of your plasmid, making the following *attB* sequence the plasmid start point:

***TTGGGCTCCCCGGGCGCGTACTCCACCTCACCCATCTGGTCCATCATGATGAACGGGTCGAGGTGG CGGTAGTTGATCCCGGCGAACGCGCGGCGCACCGGGAAGCCCTCGCCCTCGAAACCGCTGGGCGC GGTGGTCACGGTGAGCACGGGACGTGCGACGGCGTCGGCGGGTGCGGATACGCGGGGCAGCGTC AGCGGGTTCTCGACGGTCACGGCGGGCATGTCGAC***

2. Paste in entire plasmid sequence immediately following *attP* 5' boundary (below) in *pIGLETa/b* locus .gbk files (This will divide the *attP* sequence in half):

***AGCGGTTTTCGGGAGTAGTGCCCCAACTGGGGTAACCT*** **(*attB* + transgene here)**

3. Confirm 5' boundary and 3' boundary match the sequences above.
